# Into the deep: The subthalamic and para-subthalamic nuclei in behavioral avoidance

**DOI:** 10.1101/2023.07.18.549513

**Authors:** Gian Pietro Serra, Adriane Guillaumin, Bianca Vlcek, Lorena Delgado-Zabalza, Alessia Ricci, Eleonora Rubino, Sylvie Dumas, Jérôme Baufreton, François Georges, Åsa Wallén-Mackenzie

## Abstract

The subthalamic nucleus (STN) is a key component of the brain network for movement control. However, the STN is strikingly heterogeneous and also intricately engaged in limbic and cognitive functions. The STN shows aberrant firing activity in several neurological and neuropsychiatric disorders, including Parkinsońs disease (PD). Deep brain stimulation (DBS) in the STN alleviates motor impairment in PD, but patients have reported altered mood as adverse side-effect. Recent observations suggest that optogenetic STN activation in mice induces flight behavior. We hypothesized that STN activation stand at risk of causing an aversive response with behavioral avoidance as consequence. The STN is directly adjoined with the para-STN (pSTN), a hypothalamic area correlated with appetitive and aversive behavior. STN-DBS aiming to correct STN might thereby also modulate pSTN. To dissociate the impact of STN and pSTN, we took advantage of selective promoters in mice, identified in our recent RNA-sequencing of the subthalamic area, to selectively direct optogenetic excitation. Acute photostimulation resulted in aversion via both the STN and pSTN, but only STN-stimulation-paired cues resulted in conditioned avoidance. Viral-genetic tracing coupled with electrophysiological recordings identified a polysynaptic pathway from the STN to the lateral habenula, a critical hub for aversion and associated with clinical depression. This study demonstrates that STN activation is directly correlated with aversion, and thereby contributes neurobiological underpinnings to emotional affect upon STN manipulation with implications for STN-targeted treatment outcome.

---

The subthalamic nucleus (STN) is a critical component of the elaborate brain network essential for motor control ^1–8^. However, the STN is also important for associative and limbic functions ^9–16^. The STN thereby shows remarkable functional heterogeneity and its actions have broad impact on physical and mental health. Activation of the STN is correlated with motor inhibition, preventing the inititiation of unintended movement and stopping, or pausing, of ongoing movement to facilitate action selection ^7, 17–22^, functions necessary to achieve behavioral control and flexibility. Consequently, aberrant STN firing contributes to behavioral dysfunction, as observed in e.g. Parkinsońs disease (PD) ^1, 3, 23^, and obsessive-compulsive disorder (OCD) ^24^. Degeneration of STN neurons is a hallmark of supranuclear palsy and Huntingtońs disease ^25, 26^, further pinpointing the importance of STN in behavioral regulation. The STN is an important target area in deep brain stimulation (DBS), aiming to correct aberrant STN firing activity in advanced-stage PD ^9^ and highly treatment-resistant OCD ^24, 27, 28^. Based on its high success rate, STN-DBS is today a prioritized treatment of PD and OCD, as well as co-morbid PD/OCD, and has been proposed as treatment for additional neurological and neuropsychiatric disorders ^29^ such as severe and refractory Tourette’s syndrome ^30, 31^. Recent research has highlighted STN-DBS as a promising intervention candidate for substance use disorder ^32–37^. While already an important intervention target in PD and OCD, and while increasing interest is propelled towards its implementation in treatment of additional behavioral disorders, one major clinical challenge with STN-DBS that remains to solve is the emergence of disabling adverse side-effects. Some STN-DBS-induced side-effects resemble neuropsychiatric disorder, including depression ^29, 38^. Affective and cognitive side-effects draw attention to the urgency of solving the neurobiological underpinnings of STN in functions beyond motor control.

The functional heterogeneity of the STN structure has been proposed to be mediated via a tripartite functional segregation into three anatomically distinct STN subareas (also known as territories or domains) that, in turn, show distinct afferent (cortical and pallidal) and efferent (pallidal, e.g. globus pallidus *interna* (GPi)/entopeduncular nucleus (EP), ventral pallidum (VP), substantia nigra *pars reticulata* (SNr)) projectivity, forming three parallel circuits encompassing motor, associative and limbic functions, respectively ^39–42^. Further, beyond its own anatomical-functional complexity, the STN is located in an anatomical-functional heterogeneous brain area. The dorsal aspect of the STN borders to the zona incerta (ZI), another target for DBS in PD, co-morbid PD/OCD, and tremor ^41, 43, 44^, while the medial aspect of the STN, known as the “limbic tip”, is directly fused with the para-STN (pSTN), a small brain area of hypothalamic origin which was recently implicated in both appetitive and aversive responses ^45–51^. In close proximity to the medial STN and pSTN, reside the posterior and lateral hypothalamic subnuclei (PH, LH), important for feeding and homeostasis, and LH recently associated with aversion ^52, 53^. Further, the STN and pSTN are surrounded by passing fibers that spread extensively within the brain ^41, 50, 54–56^. STN-DBS treatment strategies, aiming to alter the activity of dysregulated STN neurons, thereby stand at risk of affecting functionally diverse neurons within the STN, but also anatomically neighboring structures that subserve regulation over other types of behaviors than those intended for treatment, and that instead may be the direct cause of adverse side-effects. To advance precision in treatment and reach symptom-alleviation without causing unwanted side-effects, experimental strategies in rodents that enable dissociation, and hence revelation, of the full repertoire of behaviors mediated by the subthalamic area are necessary.

In its natural brain circuitry, the STN, which is primarily excitatory and expresses the Vesicular glutamate transporter 2 (Vglut2)^57–60^, receives excitatory regulation from the cerebral cortex and inhibitory regulation from globus pallidus *externa* (GPe) ^1, 61^. Mimicking excitatory and inhibitory stimulation of the STN in an optogenetic paradigm, STN excitation causes the mice to escape the photostimulation ^58^. This finding suggests that STN excitation might cause an aversive avoidance-type behavior. Several studies have demonstrated a correlation between STN and reward ^32–37, 62^. Further, STN-DBS can induce loss of motivation ^63, 64^. However, a direct correlation between STN and aversion, providing a neurobiological framework for negative mood upon STN hyperactivation, has yet to be identified.

Here, we hypothesized that the STN exerts influence on behavioral aversion. We also hypothesized that STN-driven aversion is distinct from aversion mediated by the closely adjacent pSTN, recently implicated in aversive response. Based on our recent single-cell transcriptomics analysis within the subthalamic area, molecular markers for the STN and pSTN are available that now allow spatially precise targeting in the experimental setting in mice ^65^. Here, such promoters were implemented in optogenetics-driven neurocircuitry dissection to tease out the nature of affective behavior mediated by distinct neuronal domains within the subthalamic area.

## Results

### STN neurons express Vglut2 and Pitx2 while pSTN neurons express Tac1

The STN is located in the deep base of the caudal forebrain and is visible in a histological section as a cellular density in-between the cerebral peduncle ventrally and the ZI dorsally, and joined medially to the pSTN (Fig. 1A). In terms of molecular markers, fluorescent *in situ* hybridization (FISH) analysis of serially cryo-sectioned brains from mice at post-natal day 28 verified previous findings that the STN is positive for Vglut2 (Fig. 1B, D, F) and paired-like homedomain 2 (Pitx2) (Fig. 1C, E, F) mRNAs, with Pitx2 mRNA showing a higher level of selectivity for the STN (Fig. 1C) than Vglut2 (Fig. 1B), as Vglut2 is abundant also in surrounding thalamic and hypothalamic structures (Fig. 1B-C). Within the STN, Pitx2 and Vglut2 show 100% overlap ^66^, and STN^Pitx2/Vglut2^ double-labeling covers the rostro-caudal extent of the elongated STN structure (Fig. 1D-F). However, while present throughout the STN, Vglut2 mRNA is somewhat denser in the medial than lateral STN, creating a medial-lateral Vglut2 mRNA gradient (Fig. 1D, compare D1, D2). Further, parvalbumin (PV), a marker for a subset of STN neurons, shows denser distribution dorsally than medially, forming an opposite gradient (Fig. 1G-I), in accordance with previous literature (STN^PV^) ^65, 67–70^.

**Figure 1.**
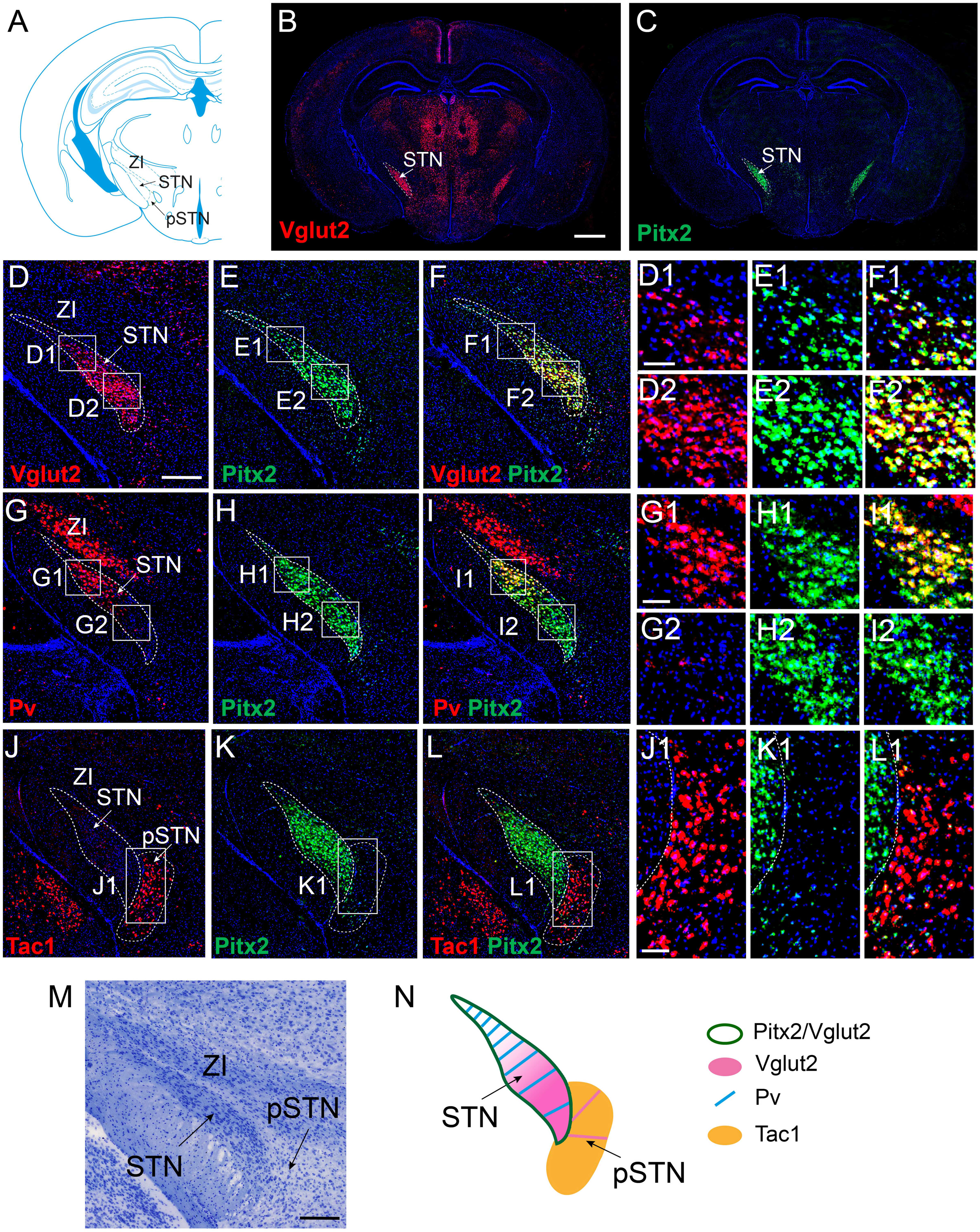
Distinct STN neurons positive for Pitx2, Vglut2 or PV while pSTN neurons positive for Tac1. (A-N) Histological analysis on coronal mouse brain sections of mRNAs representing whole STN (Pitx2, Vglut2), subtype STN neurons (PV), or pSTN (Tac1). (A) Schematic illustration of mouse coronal section including subthalamic nucleus (STN), para- subthalamic nucleus (pSTN), zona incerta (ZI) at bregma -2.18 ^145^. B-L) Fluorescent *in situ* hybridization (FISH); detection of mRNA in coronal brain sections at level shown in (A) from postnatal day (P) 28 mice: B-C) overviews, Vglut2 (B), Pitx2 (C); D-L) close- ups of STN/pSTN/ZI area with squares pointing to cellular close-ups displayed in F1- L1 panel to right, Vglut2 (D-D2), Pitx2 (E-E2), (Vglut2/Pitx2)(F-F2); PV (G-G2), Pitx2 (H-H2), PV/Pitx2 (I-I2); Tac1 (J-J1), Pitx2 (K-K21), Tac1/Pitx2 (L-L1). N) Histological staining (cresyl violet) of same area as in A-M, arrows pointing to STN, pSTN, ZI. O) Schematic summary of mRNA patterns in STN and pSTN as identified in FISH analysis above. Serial sections throughout the STN/pSTN area were analysed for each probe pair in number of mouse brains precised as: Vglut2 (n=8 brains), Pitx2 (n=8 brains), PV (n=5 brains), Tac1 (n=4 brains). Scale bars: B: 1 mm; D: 300 µm; D1, H1 and K1: 70µm, M: 300 µm.

Some Pitx2^+^ and Vglut2^+^ cells can be found in the pSTN (Fig. 1D-F) whereas ZI, an inhibitory structure, is negative for both Pitx2 and Vglut2 mRNAs (Fig 1D-F). In our recent single nuclei (sn)RNAseq analysis of the subthalamic area, Pitx2, Vglut2 and PV were all identified and subsequently mapped to the STN with histological FISH analysis ^65^. In addition, tachykinin precursor 1 (Tac1) was identified, and FISH analysis demonstrated Tac1 mRNA as a selective marker for pSTN (pSTN^Tac1^) ^65^. In accordance, prominent Tac1 mRNA was here detected in the pSTN, with very few positive cells in the STN (Fig. 1J-L). A majority of Tac1^+^ cells in the pSTN were negative for Pitx2. Summarizing the results obtained in histological analysis of the STN and pSTN, Pitx2 and Vglut2 mRNAs both represent the whole extent of the STN, with a medial-lateral gradient of Vglut2, PV represents abundant, but scattered, neurons across the STN in a lateral-medial gradient, and Tac1 mRNA represents the pSTN, with only the occasional Tac1^+^ neuron detected in the STN (Fig. 1M, N).

### Optogenetic stimulation of STN and pSTN causes different avoidance-type behaviors

Taking advantage of *Pitx2*, *Vglut2*, *PV* and *Tac1* promoter activities, the corresponding Cre-recombinase transgenic mouse lines were used as drivers of floxed alleles to allow optogenetics-driven circuitry dissection segregating the STN and pSTN (whole-STN, Vglut2^Cre^ and Pitx2^Cre^; STN PV^+^ subpopulation, PV^Cre^; pSTN, Tac1^Cre^). Cre-mice (also referred to as recombinase mice) were injected bilaterally with adeno-associated (AAV) virus to express a construct encoding either Channelrhodopsin-2 (ChR2) fused with a fluorescent reporter gene (enhanced yellow fluorescent protein (eYFP) construct) encoding YFP (experimental groups, abbreviations: STN^Pitx2^/ChR2, STN^Vglut2^/ChR2, STN^PV^/ChR2, pSTN^Tac1^/ChR2) or, as controls, a similar construct encoding YFP but lacking ChR2 (control groups, abbreviations: STN^Pitx2^/CTRL, STN^Vglut2^/CTRL, STN^PV^/CTRL, pSTN^Tac1^/CTRL)(Fig. 2A). Fiber optic probes were placed bilaterally above the STN (Fig. 2A). Verification of optical probe position and distribution of YFP was performed in histological analysis on brain sections upon completion of functional analysis (electrophysiology, behavior) which confirmed similar distribution of each respective promoter activity as corresponding mRNA (Pitx2^mRNA^/Pitx2^Cre^; Vglut2^mRNA^/Vglut2^Cre^; PV^mRNA^/PV^Cre^; Tac1^mRNA^/Tac1^Cre^)(Fig. 2B-E, Suppl. Fig. 1).

**Figure 2.**
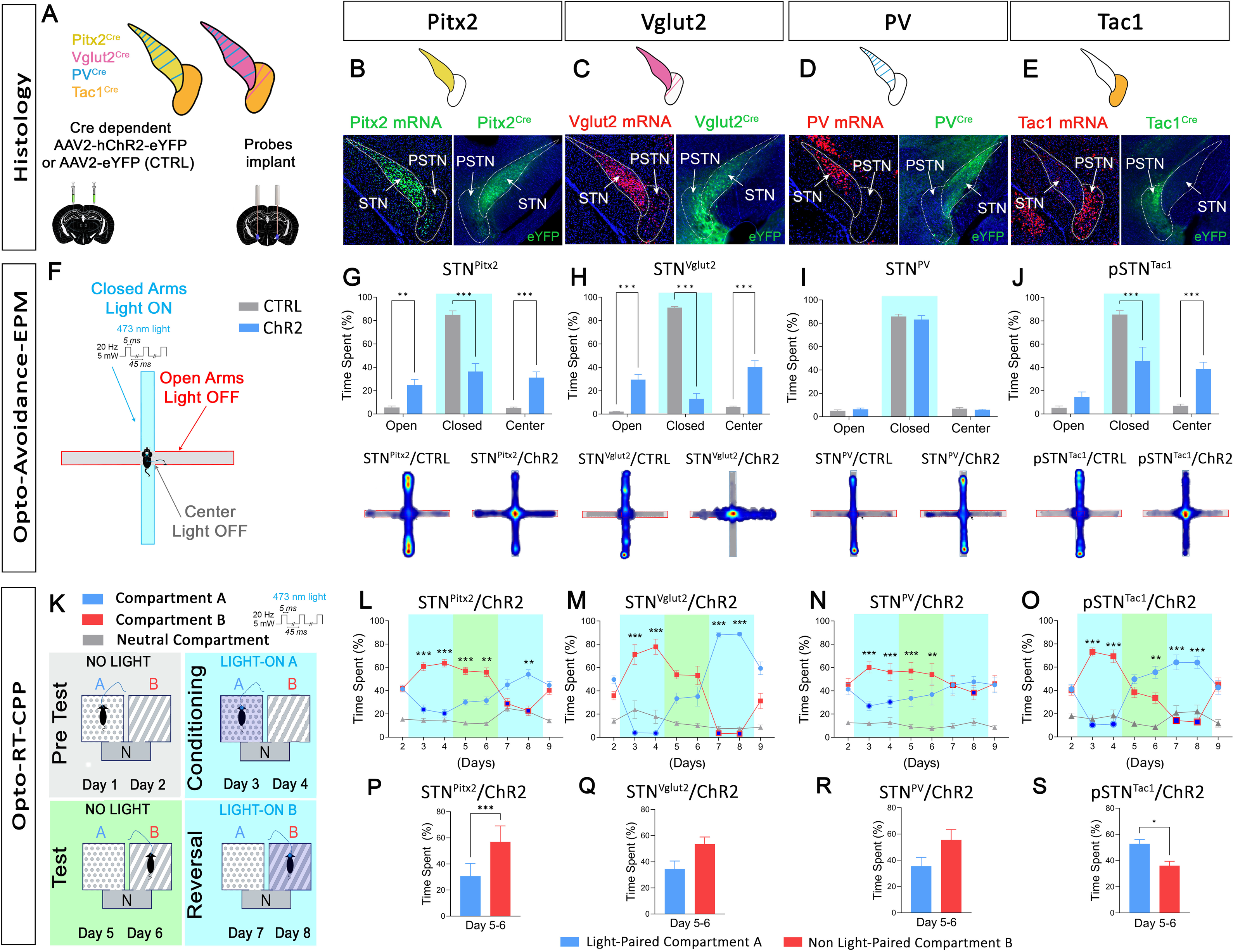
Optogenetic stimulation of STN and pSTN causes different avoidance- type behaviors. A) Graphical illustration. Pitx2^Cre^, Vglut2^Cre^, PV^Cre^, Tac1^Cre^ transgenic mice (representing STN or pSTN; Fig. 1) were stereotaxically injected into STN or pSTN with rAAV carrying Cre-dependent ChR2-YFP construct (ChR2) or YFP construct alone (control; CTRL) and fiber optic probes were implanted bi-laterally above the STN/pSTN area. B-E) Histological analysis on coronal sections from injected mice displaying each mRNA (left, FISH) and corresponding Cre-driven eYFP (right, immunofluorescence). Pitx2/Pitx2^Cre^ (B), Vglut2Vglut2^Cre^ (C), Pv/PV^Cre^ (D), Tac1/Tac1^Cre^ (E). Suppl. Fig. 1 supporting data. F-J) Opto-Avoidance-EPM test. Graphical illustration of setup (F), Time spent in arms (G-J): CTRL vs. ChR2 **p<0.01, ***p<0.001, and below, representative heat map: STN^Pitx2^/ChR2 (n=15) and STN^Pitx2^/CTRL mice (n=14) (G), STN^Vglut2^/ChR2 (n=7) and STN^Vglut2^/CTRL (n=6) (H), STN^PV^/ChR2 (n=9) and STN^PV^/CTRL (n=11) (I), pSTN^Tac1^/ChR2 (n=11) and pSTN^Tac1^/CTRL (n=12) (J) mice. K-S) Opto-RT-CPP test. Graphical illustration of setup. Blue, Compartment A, Red, Compartment B, Gray, Neutral compartment (K). (L-O) Percentage of time spent. Compartment A *vs*. Compartment B, **p<0.01, ***p<0.001, and (P-S) Averaged percentage of time (days 5-6) spent in each compartment: light-paired *vs.* non-light-paired, *p<0.05, ***p<0.001: STN^Pitx2^/ChR2 (n=15) (L,P),M) STN^Vglut2^/ChR2 (n=7) (M,Q), STN^PV^/ChR2 (n=9) (N,R), pSTN^Tac1^/ChR2 (n=11) (O,S) mice. Blue, Light-paired Compartment A, Red, non-light-paired Compartment B, Dark blue filled circles (Compartment A) and squares (Compartment B) indicate photostimulation (Light-ON). (n) refers to number of mice. Suppl. Fig 3, 4 and Suppl File 1, supporting data. Data expressed as mean ±SEM. Abbreviation: FISH; Fluorescent *in situ* hybridization.

To first validate the stimulation protocol, a series of electrophysiological analyses were performed in STN^Pitx2^/ChR2 mice. *In vivo* single cell electrophysiological recordings upon optogenetic stimulation of the STN were performed (Suppl. Fig. 2). An optotagging protocol (Suppl. Fig. 2) was used to stimulate and record within the STN. To observe the reaction of STN neurons to photostimulation, peri-stimulus time histograms (PSTH protocol, 0.5 Hz, 5 ms bin width, 5-8 mW) were created by applying a 0.5 Hz stimulation protocol for at least 100 seconds. Action potentials in ChR2- positive STN cells were successfully evoked by STN photostimulation (Suppl. Fig. 2). Once neuronal activity of STN neurons returned to baseline, a photostimulation protocol intended for behavioral experiments was validated (Behavior protocol, 20 Hz, 5 ms pulses, 5-8 mW; 10 seconds) which increased the frequency and firing rate of STN neurons for the whole duration of the stimulation, after which they returned to normal (Suppl. Fig. 2).

The excitability of STN neurons was next confirmed by patch-clamp recordings in the STN of STN^Pitx2^/ChR2 mice with the optic probe placed above the recording site (Suppl. Fig. 2). ChR2-YFP expression was strong in the STN (Suppl. Fig. 2), and all of the STN neurons tested responded to continuous (Suppl. Fig. 2) or 20 Hz trains (Suppl. Fig. 2) of light stimulation by sustained ChR2-mediated currents. When light stimulation was applied in brain slices from STN^Pitx2^/CTRL control mice, no current was observed in STN neurons.

A series of behavioral analyses was next performed to identify and dissociate the nature of affective behavior induced upon optogenetic stimulations within the STN and pSTN by comparing all four recombinase mouse lines (Suppl. File 1, detailed behavior statistics). First, based on previous reports of grooming and jumping behavior upon STN stimulation ^58, 71^, mice were tested in an open field apparatus coupled with laser activation (Opto-OpenField) (Suppl. Fig. 3). Behavior during periods of photostimulation (Light-ON) was compared with periods of no photostimulation (Light-OFF). Natural grooming follows a cephalo-caudal rule and is a normal cleansing behavior for rodents ^72, 73^. However, when accentuated above or below baseline, rodent self-grooming has been correlated with anxiety and/or aversion, revealing signs of distress ^74, 75^. Indeed, also certain compulsions in humans, such as excessive self-grooming, have been suggested to manifest behavioral repetitions in attempts to relieve anxiety and/or aversion caused by strongly unpleasant sensations ^76^. STN- induced grooming has been described as strongly repetitive and directed at the face region only ^58^. Confirming previous reports, STN stimulation caused repetitive face grooming immediately upon start of photostimulation (Light-ON) (Suppl. Fig. 3). This was observed for all STN^Pitx2^/ChR2 mice, some of which also showed escaping jumping behavior. The other STN-cohort mice, STN^Vglut2^/ChR2, responded to photostimulation to such a substantial extent that excessive jumping behavior precluded grooming analysis. On the other hand, STN^PV^/ChR2 mice, representing a subpopulation STN neurons, did not respond to photostimulation with any signs of jumping or grooming, instead this group of mice was similar to all control groups (Suppl. Fig. 3). When addressing the pSTN, mice did not jump or escape the arena upon stimulation, but they did perform repetitive face grooming. pSTN^Tac1^/ChR2 mice showed a different response curve than STN^Pitx2^/ChR2 mice, as the grooming did not stop immediately as photostimulation stopped (Suppl. Fig. 3). Thus, STN and pSTN recombinase mice showed distinctly different grooming phenotypes.

Taking this result into consideration, a new protocol was designed to address optogenetic-induced place avoidance in the standardized elevated plus maze (EPM) apparatus. The EPM consists of four arms elevated from the floor and connected via a center area. Two arms are open and exposed, providing a naturally aversive space that mice avoid to spend time in. The two other arms are enclosed, providing a sheltered space than mice prefer over the open and naturally aversive arms. The center area is less exposed than the open arms, but not sheltered. In a standard EPM test, mice prefer the closed arms and avoid the open arms and center, thus showing place avoidance to the naturally aversive space ^77, 78^. In this new protocol, referred to as the Opto-Avoidance-EPM test, entry into either of the closed, sheltered arms is paired with subthalamic photostimulation (Light-ON) (Fig. 2F). Entry into either of the center or open arms remains unpaired with photostimulation (Light-OFF). Thus, by pairing the sheltered area with optogenetic activation, the Opto-Avoidance-EPM protocol allows direct comparison between a naturally aversive context (open arm) and aversion caused by subthalamic activation (closed arm). Applying this new protocol, both STN (Pitx2 and Vglut2) (Fig 2G,H, Suppl. Fig. 4) and pSTN (Tac1) (Fig 2J, Suppl. Fig. 4) recombinase mice actively avoided the photostimulation-paired closed arms and instead showed increased number of entries and time spent in the photostimulation-unpaired, but naturally aversive, open arms. Further, also the center area, photostimulation-unpaired and more sheltered than the exposed open arms, gained in visits. In contrast, none of the control groups, nor STN^PV^/ChR2 mice, showed any effect of photostimulation, instead they all preferred the closed arms, as in a standard EPM test, despite the exposure to photostimulation (Fig. 2G-J; Suppl. Fig. 4). Analysis in the Opto-Avoidance-EPM test thereby demonstrated direct causality between activation within different locations within the subthalamic area (STN and pSTN) and behavioral avoidance. Further, the avoidance response was significantly more pronounced than that to natural aversion, correlating the observed behavior with a strongly aversive response. While a similar response was observed during both STN and pSTN stimulation, it was also evident that STN-induced aversion was not mediated by the STN PV^+^ subpopulation.

Based on these findings, to further dissociate the nature of behavioral avoidance observed upon STN *vs* pSTN stimulations, a new protocol to address optogenetically-induced aversion using the standardized conditioned place preference (CPP) apparatus was designed. In a standard CPP, two compartments (A and B) that differ in terms of contextual cues, are connected via a transparent corridor, and preference *vs* avoidance to either compartment upon presentation various types of stimuli, and stimuli-paired cues, can be assessed. Here, to allow analysis of both immediate (real-time) and long-term effects, a protocol here referred to as Opto-RT-CPP was designed to address response to photostimulation both in real-time (RT) and upon conditioning to photostimulation-paired cues, and finishing off with reversal of photostimulation (Fig. 2K). Following habituation (Day 1) and pre-test day (no light, Day 2) to reveal any bias, entry into compartment A was paired with subthalamic photostimulation (Light-ON A, Day 3,4) after which followed two test days of conditioned response (no light, Day 5,6). After this test followed a reversal test to assess ability to reverse the learned response in which compartment B, instead of A, was paired with photostimulation (Light-ON B, Day 7,8) and one test day (no light, Day 9) (Fig. 2K). As expected, none of the control groups showed photostimulation-related avoidance at any time (Suppl. Fig. 4). In contrast, STN mice (STN^Pitx2^/ChR2, STN^Vglut2^/ChR2) actively avoided any chamber paired with photostimulation. All STN mice spent significantly less time in the photostimulation-paired compartment than the unpaired compartment when assessed in real-time (Fig. 2L, M). STN mice also demonstrated conditioned avoidance on test days, but only STN^Pitx2^/ChR2 reached statistical significance during test days (Fig. 2L, M, P, Q).

Next, when addressing the PV-subpopulation of STN neurons, STN^PV^/ChR2 mice showed a similar response as STN^Pitx2^/ChR2 mice both during real-time stimulation and when testing for a conditioned response, but did not show reversal (Fig. 2N, R).

Finally, when assessing how mice behaved upon pSTN stimulation, an opposite type behavior was revealed, at least in terms of conditioning (Fig. 2O, S). This was unexpected, given that a recent study used a similar (not identical) place-preference setup and could show aversion upon acute optogenetic stimulation of pSTN using Vglut2^Cre^ mice ^51^. In the extended protocol implemented here, pSTN^Tac1^/ChR2 mice showed real-time place aversion, but failed to show conditioned place avoidance. Instead, when addressing context-conditioned response, it was evident that pSTN-activation caused conditioned preference, rather than avoidance, to the photostimulation-paired compartment (Fig. 2O, S). Similar as STN mice, pSTN mice showed reversal with the compartmental change in source of stimulation, but none of the groups reached significance on the reversal test day.

Taken together, these analyses identified a robust correlation between STN activation and place avoidance both upon direct stimulation (Opto-Avoidance-EPM and Opto-RT- CPP; real-time) and upon exposure to aversion-paired cues (Opto-RT-CPP; test days). While STN and pSTN mice all responded with grooming in the Opto-OpenField, pSTN mice did not show either escaping behavior or conditioned place avoidance. Instead pSTN activation caused the opposite, conditioned place preference. This battery of tests thereby allowed the functional dissociation between the anatomically adjacent pSTN and STN in the context of conditioned approach (pSTN) *vs* avoidance (STN) behavior.

### STN stimulation, but not pSTN stimulation, suppresses sugar positive reinforcement

Based on the findings above, we hypothesized that optogenetic stimulation causing aversive behavioral avoidance should be sufficient to induce a negative reinforcement behavior. To test this hypothesis, we next attempted to assess whether STN^Pitx2^/ChR2 mice could learn to make active nose-pokes to terminate STN optogenetic activation. In this opto-negative reinforcement paradigm (Opto-NR), all STN^Pitx2^/ChR2 mice failed to acquire an active nose-poke response to terminate optogenetic stimulation and the experiment was aborted.

Instead, we decided to address the hypothesis that STN optogenetic activation, shown to induce place avoidance, would disrupt positive reinforcement behavior. We also hypothesized that STN and pSTN mice would respond differently, given their differential response in the Opto-RT-CPP conditioning test. To test these hypotheses, an instrumental sugar positive reinforcement paradigm (Sugar-PR) was implemented using operant boxes equipped with two nose-poke (NP) apertures coupled to a reward dispenser with one active setup (NP leads to sugar delivery; active NP) and one inactive setup (no sugar delivery; inactive NP). The original schedule applied consisted of 3 phases (Phase 1, training to nose-poke for sugar reward in a fixed ratio (FR) paradigm; Phase 2, nose-poke to earn sucrose paired with subthalamic photostimulation, Phase 3, sugar reinstatement; photostimulation removed (Fig. 3A, and Suppl. Fig. 5).

**Figure 3.**
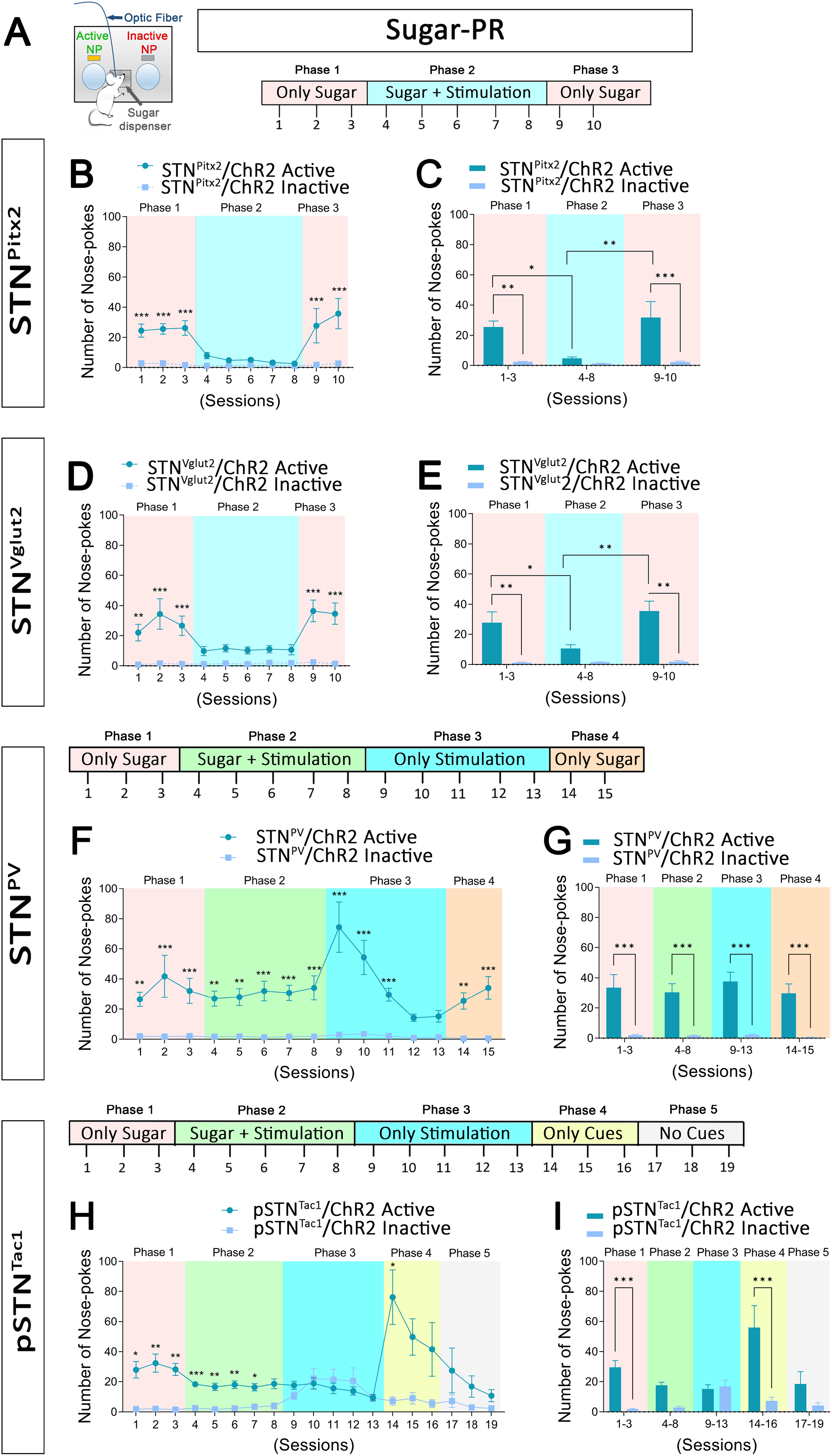
STN stimulation, but not pSTN stimulation, suppresses sugar positive reinforcement. A-I) Positive reinforcement (PR) self-stimulation test. Graphical illustration of setup and original protocol (A). B-C) Results, STN^Pitx2^/ChR2 mice (n=8). Number of active *vs.* inactive nose-pokes: ***p<0.001 (B). Average number of active and inactive nose- pokes/phase: phase 1 inactive *vs.* active, **p<0.01; phase 3 inactive *vs.* active ***p<0.001; active nose-pokes phase 1 *vs.* phase 2, *p<0.05; active nose-pokes phase 2 *vs.* phase 3, **p<0.01 (C). D-E) Results, STN^Vglut2^/ChR2 mice (n=5). Number of STN^Vglut2^/ChR2 active *vs.* inactive nose-pokes: **p<0.01, ***p<0.001 (D). Average number of STN^Vglut2^/ChR2 active and inactive nose-pokes/phase: phase 1 inactive *vs.* active, **p<0.01, phase 3 inactive *vs.* active **p<0.01; active nose-pokes phase 1 *vs.* phase 2, *p<0.05; active nose-pokes phase 2 *vs.* phase 3, **p<0.01 (E). F-G) Protocol and results, STN^PV^/ChR2 mice (n=9). Number of active *vs.* inactive nose-pokes: **p<0.01, ***p<0.001 (F). Average number of STN^PV^/ChR2 active and inactive nose- pokes/phase: inactive *vs.* active, ***p<0.001 (G). H-I) Protocol and results, pSTN^Tac1^/ChR2 mice (n=11). Number of active *vs.* inactive nose-pokes: *p<0.05, **p<0.01, ***p<0.001 (H). Average number of pSTN^Tac1^/ChR2 active *vs.* inactive nose- pokes/phase: ***p<0.001 (I). (n) refers to number of mice. Suppl. Fig 5, 6 and and Suppl File 1, supporting data. Data expressed as mean ±SEM.

In the absence of photostimulation (Phase 1), all STN^Pitx2^/ChR2 (Fig. 3B, C) and control (Suppl. Fig. 6) mice showed a significant higher number of active compared to inactive NPs, earning sugar reinforcers. This was the expected response, given the positively reinforcing properties of sugar. With the coupling to STN-photostimulation (Phase 2), control mice continued the same behavior whereas STN^Pitx2^/ChR2 mice responded strongly with reduced nose-poke activity on the active NP, a direct consequence of optogenetic STN activation. The difference between active and inactive NPs was no longer significant as the active NPs dropped with the onset of photostimulation. With subsequent removal of photostimulation (Phase 3), STN^Pitx2^/ChR2 mice resumed nose-poking (similar to Phase 1), with the number of active NPs significantly higher than the inactive ones. No difference in the number of active NPs were observed between STN^Pitx2^/ChR2 and control mice during Phases 1 and 3, while there was a significant decrease selectively for STN^Pitx2^/ChR2 mice in the photostimulation phase (Suppl. Fig. 6). Similar results were obtained with STN^Vglut2^/ChR2 mice, which made more active than inactive nose-pokes during phases 1 and 3, but with significant decrease during phase 2, leading to significantly lower number of active NPs in phase 2 compared to both phase 1 and 3 (Fig. 3D, E).

When testing the STN^PV^/ChR2 mice, no effect of STN-photostimulation was observed. (Fig. 3F, G). To address this result further, the previous sugar reinstatement phase (Phase 3) was changed for optogenetic stimulation only (instead of sugar only), administered as a consequence of the active NPs (Suppl. Fig. 5). This led to an initial increase in the number of active NPs (seeking) due to the absence of the reinforcing stimulus (sugar), followed by a progressive decrease in number of active NPs (extinction). After this followed a Phase 4, in which sugar was again available (Fig. 3F, G). As expected, this restoration of sugar only caused the number of active NPs to again become significantly larger than the inactive ones. Thus, the mice responded to sugar but not to STN photostimulation; no difference was observed between STN^PV^/ChR2 and control mice in any of the 4 phases (Suppl. Fig. 6). This result supports the outcome of the Opto-Avoidance-EPM test in that the PV^+^ STN subpopulation does not play a first order role in aversive response.

The effect of pSTN stimulation in the positive reinforcement test resulted in a response that did not resemble aversion. pSTN^Tac1^/ChR2 mice did not show a significant aversive response upon optogenetic stimulation. Unexpectedly, pSTN mice continued to nose-poke for sugar in the presence of photostimulation (Fig. 3H, I). Further, using a Phase 3 in which active NPs were coupled with photostimulation alone (no sugar), no decrease in the number of active NPs was achieved, but instead the number of inactive NPs increased until reaching the same values as the active ones (Fig. 3H, I). To assess this surprising response further, two subsequent phases were added. Now, pSTN-photostimulation and sugar delivery were removed, leaving the acoustic cue as consequence of the active NPs (Phase 4), and subsequently removing also the cue, so that active NPs were equal to inactive NPs (Phase 5) (Suppl. Fig. 5). pSTN^Tac1^/ChR2 mice responded during the first sessions of Phase 4 with strongly increased number of active NPs (seeking) and then proceeded towards extinction during the following sessions (Fig. 3H). This result was different to what observed with controls as these showed a strong increase in active NPs when sugar was removed (Suppl. Fig. 6). In contrast to STN-mediated aversion, optogenetic activation of the pSTN did not negatively affect sugar consumption, instead mice maintained their self-administration activity in the presence of pSTN stimulation.

### Distinctly different projections for STN and pSTN

Projection targets of STN^Pitx2^, STN^Vglut2^, STN^PV^ and pSTN^Tac1^ neurons were next compared to identify circuitry components of the identified behavior (Fig. 4). Distribution patterns of the YFP reporter was analyzed by immunofluorescent analysis of STN and pSTN neurons and their projections on serial sections derived from injected recombinase mice (Fig. 4A). YFP-labeling in projections from STN^Pitx2^ neurons reached pallidal structures, the EP (the rodent counterpart of primate GPi), GP (the rodent counterpart of primate GPe), and VP, and also reached nigral structures (SNr and substantia nigra *pars compacta*, SNc) (Fig. 4B) in accordance with our recent report^58^. STN^Vglut2^ and STN^PV^ neurons showed similar, but not identifical, projections (Fig. 4C, D). STN^Pitx2^ and STN^PV^ projections were strongest in EP, GP, SNr (Fig. 4B, D). In contrast, pSTN^Tac1^ neurons primarily projected to central amygdala, bed nucleus of stria terminalis (BNST), and both medial and lateral septum, areas associated with limbic functions, while pSTN^Tac1^ projections to GP and SNr were sparse or non-existant (Fig. 4E). STN^Vglut2^ also showed projections to BNST, central amygdala and septal nuclei (Fig. 4C), whereas STN^PV^ showed no YFP labeling in these areas and where STN^Pitx2^ projections were either weak or absent. Further, no STN^Pitx2^ projections were observed in the LHb.

**Figure 4.**
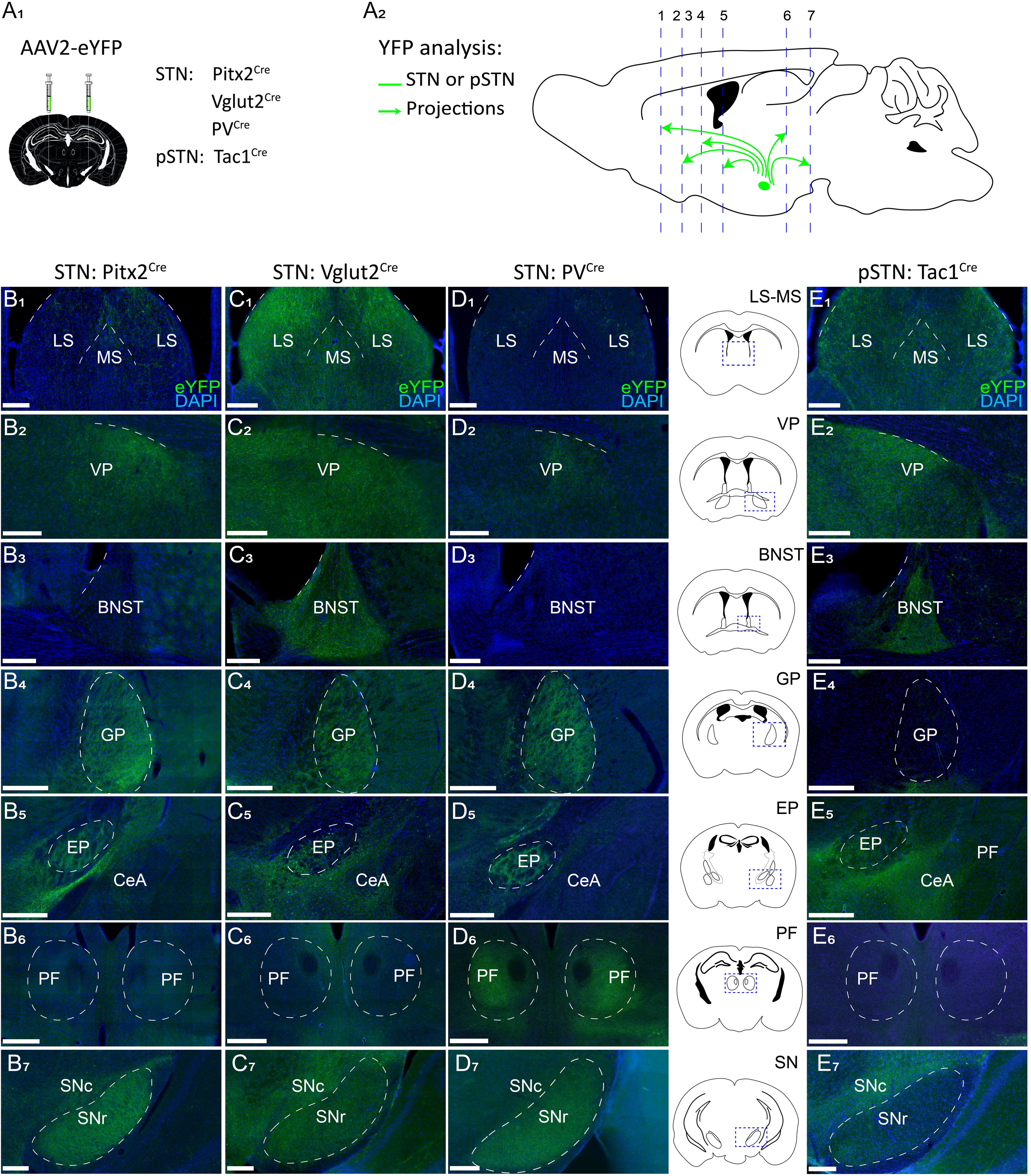
Distinctly different projection patterns for STN and pSTN. A-D) Projection analysis in brains from STN^Pitx2^ (B), STN^Vglut2^ (C), STN^PV^ (D), pSTN^Tac1^ (E) mice injected with AAV2-eYFP. Viral injection strategy (A1). Illustration of sagittal mouse brain indicating with dotted lines the seven section levels displayed as coronal sections below (A2). eYFP immunofluorescence detecting STN (B-D) or pSTN (E)- derived projections (in green) of STN^Pitx2^ (n=29)(B1-B7), STN^Vglut2^ (n=29)(C1-C7), STN^PV^ (n=18)(D1-D7) and pSTN^Tac1^ (n=18)(E1-E7) mice. Panel between D and E; illustrations of areas shown per section level (square). (n) refers to number of mice. Representative examples displayed. Abbreviations: VP, ventral pallidum; GP: globus pallidus; EP: entopeduncular nucleus; CeA: central amygdala; BNST: bed nucleus of the stria terminalis; PF: parafascicular thalamic nucleus; SNr: substantia nigra *pars reticulata*; SNc: substantia nigra *pars compacta*; LS: lateral septum; MS: medial septum. Suppl. Fig. 1 supporting data; STN and pSTN, injected brain areas.

Since excitation of the STN, but not the pSTN, promoted both active and conditioned avoidance, sufficiently potent to suppress sugar self-administration, exploring such a putative STN neurocircuitry of aversion was of particular interest.

### Optogenetic stimulation of STN is sufficient to induce excitatory post-synaptic responses in LHb

STN-DBS has repeatedly been reported to give rise to depression as adverse side- effect ^29, 38^, providing a clinical association between STN stimulation and negative emotional affect. Furthermore, STN-HFS in rodents induces c-Fos expression in LHb and alters LHb neuron activity ^79, 80^, suggestive of functional connectivity between STN and the LHb. However, our projection analysis did not reveal any YFP-positive projections in the LHb of STN^Pitx2^ mice. Instead, STN^Pitx2^/ChR2 mice showed abundant projections to pallidal structures VP and EP, as verified above and previously reported^58^. Both VP and EP, in turn, have been shown to project to LHb ^81–84^.

To first explore any potential impact of STN-activation on LHb activity, *in vivo* extracellular recordings were performed in STN^Pitx2^/ChR2 mice (Fig. 5A). An optic fiber was positioned above the STN and a stimulation protocol (PSTH 0.5 Hz, 5 ms pulses, 5-8 mW; at least 100 seconds) was implemented (Fig. 5B). Post-recording, pontamine staining and neurobiotin-marked neurons confirmed the positioning of the recording electrodes within the LHb (Fig. 5C, D). Analysis showed that STN stimulation evoked an excitatory response in 50% of the recorded LHb neurons (onset latency =10.08 ms +/- 0.81 ms) (Fig. 5E, F). Thus, a functional STN – LHb connection could be observed. With an apparent lack of direct connection between the STN and the LHb in STN^Pitx2^ mice (confirmed by the absence of eYFP fibers in LHb; Fig. 5D), identification of the putative pathway responsible for the STN-induced post-synaptic responses in LHb was next approached.

**Figure 5.**
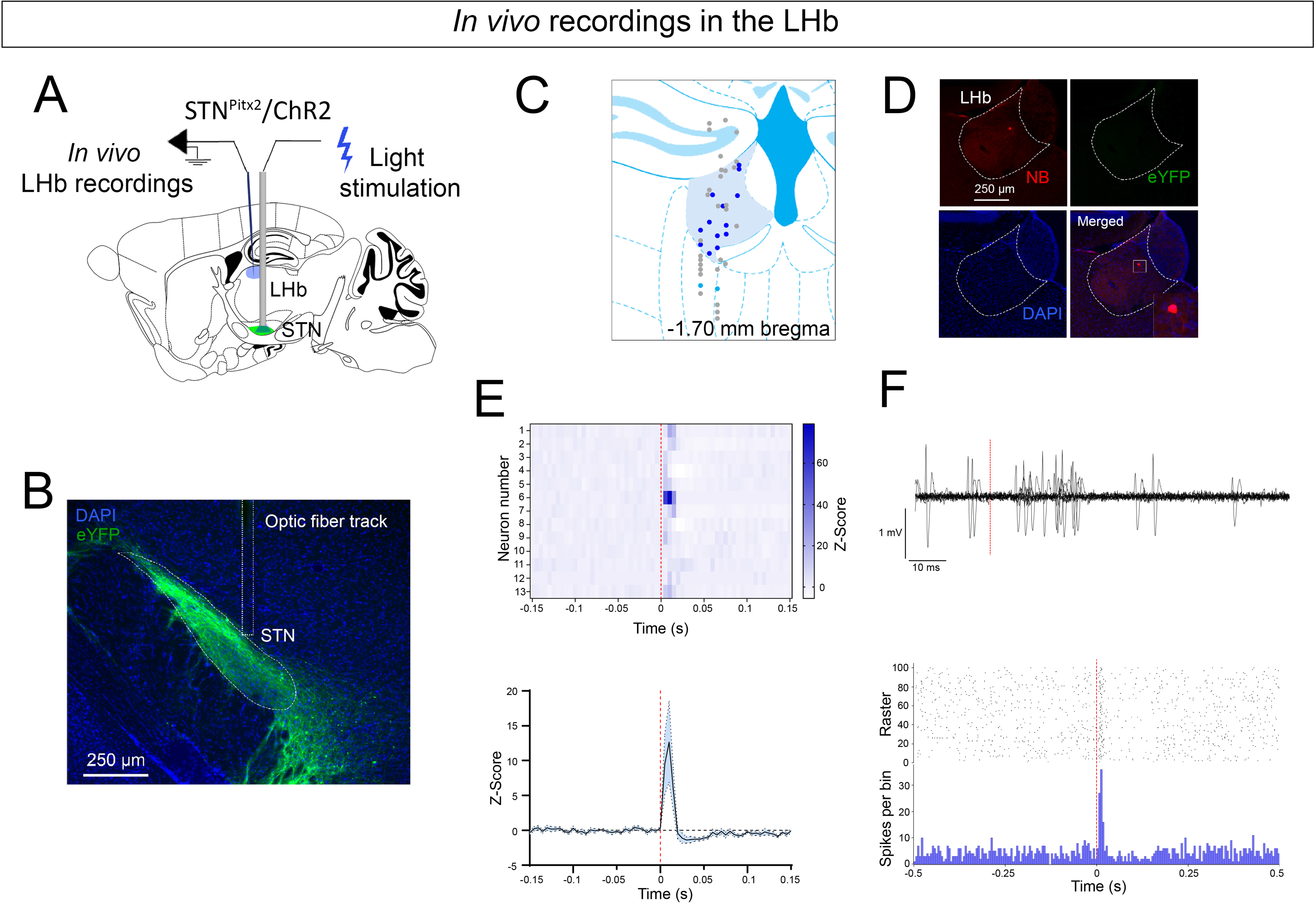
**Photostimulation in STN^Pitx^**^2^**/ChR2 mice excites STN neurons and induces excitatory post-synaptic responses in neurons of the lateral habenula (LHb)** A) LHb recording procedure with STN-photostimulation in STN^Pitx2^/ChR2 mice. B) Representative coronal image showing YFP fluorescence (green) in STN^Pitx2^/ChR2 mice with optic cannula track above the STN; nuclear marker (DAPI, blue); scale, 250 µm. C) Reconstructed mapping of LHb recorded neurons (N=44 neurons recorded in 3 mice). Dark blue dots represent LHb neurons excited during stimulation; light blue dots correspond to excited neurons outside of the LHb; grey dots correspond to not responding neurons. D) Excited LHb neuron injected with neurobiotin. No YFP-positive fibers in the LHb. E) PSTH heatmap of excited LHb neurons upon STN- photostimulation of the STN (top) and average of normalized PSTH (bottom; n=13 neurons). F) Overlap of 10 traces of a LHb neuron responding to STN photostimulation centred on the 5 ms light pulse (top) and PSTH and raster of a representative excited LHb neuron (below).

### Selective optogenetic activation of the STN reveals two di-synatic circuits connecting the STN and the LHb

Based on current literature, the STN-VP and STN-EP pathways were next examined in the context of projections to the LHb. Using rgAAV2-DIO-eYFP injections in the LHb of Vglut2^Cre^ mice, we retrogradely labeled Vglut2-expressing (glutamatergic) LHb-projecting VP and EP neurons. An AAV5-DIO-CHR2-mCherry vector was also injected in the STN to selectively manipulate STN-VP and STN-EP pathways (Fig. 6A, B). Two-three weeks later, acute brain slices were prepared and eYFP-expressing VP and EP neurons were recorded in voltage-clamp configuration and filled with biocytin (Fig. 6C- D, E,H). STN inputs were optically-stimulated and synaptic inputs were recorded in eYFP-positive VP and EP neurons, respectively. 100% of the recorded LHb-projecting EP were identified as responsive to STN inputs stimulation, while only 50% of LHb-projecting VP display light-evoked synaptic currents (Fig. 6F-G; I-J). The amplitude of the excitatory post-synaptic currents (EPSC) was significantly greater (Fig. 6K) and the latency of the EPSC significantly shorter (Fig. 6L) in EP neurons compared to VP neurons. These results suggest that two distinct di-synaptic pathways relay the STN to the LHb. Based on these results, and the fact that the anatomical proximity between the EP and the STN makes it impossible to specifically optogenetically manipulate STN terminals in the EP without directly affecting the STN, the impact on affective behavior upon selective stimulation of STN terminals in the VP was next assessed.

**Figure 6.**
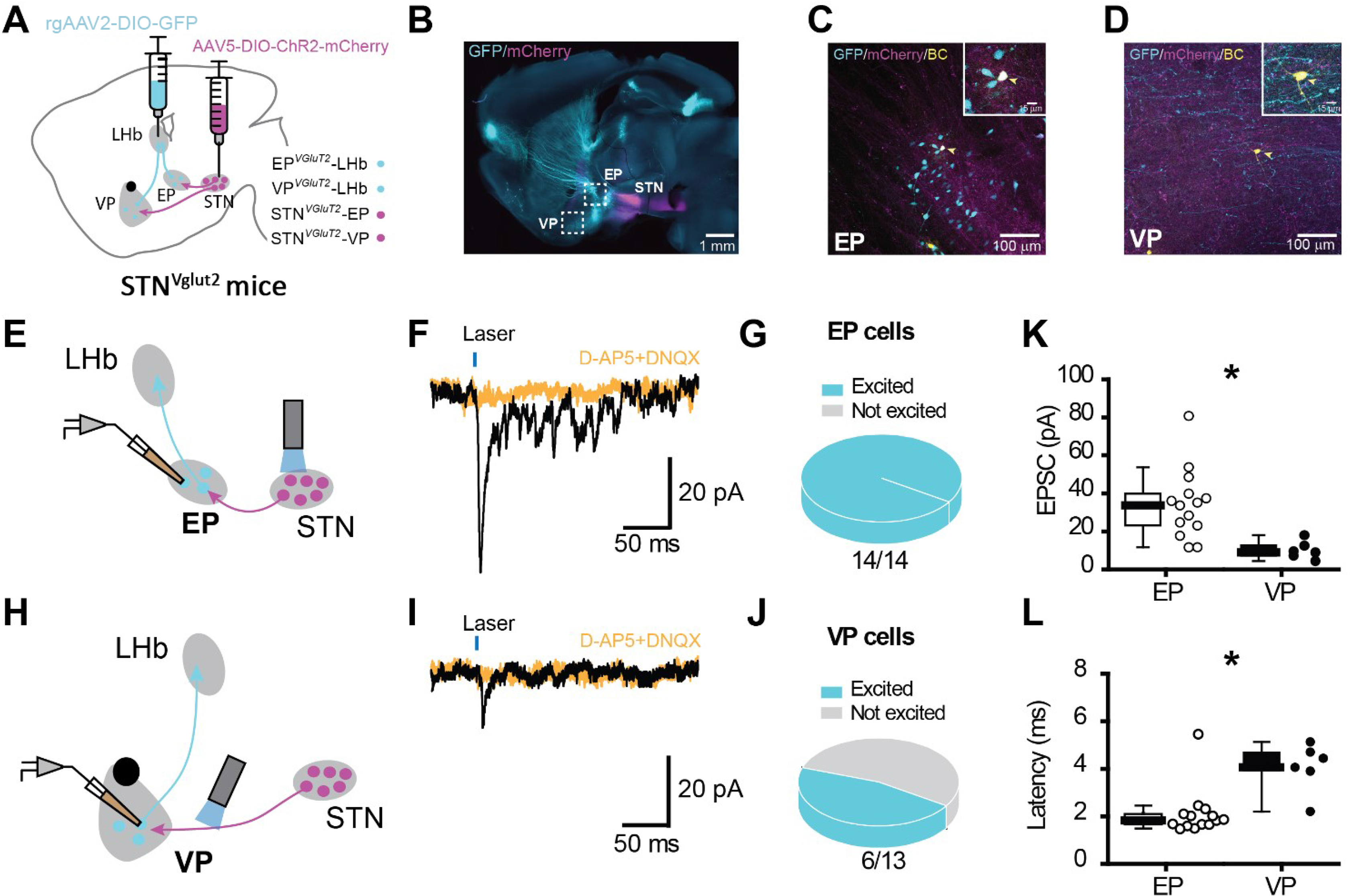
Optogenetic activation of STN projections induces excitatory post- synaptic responses in both VP and EP neurons projecting to the lateral habenula (LHb) A) Viral tracing strategy combining retro adeno-associated virus (rgAAV2-DIO-GFP) into the lateral habenula (LHb) along with a AAV5-DIO-ChR2-mCherry virus into the STN of Vglut2^Cre^ mice (STN^Vglut2^). B) Sagittal section showing mCherry (magenta) and GFP (cyan) labelling 4 weeks after the injections. Confocal images with higher magnification (upper quadrants) from the EP (C) and the VP (D) depecting in vitro recorded neurons filled with the BC (arrowheads) and expressing the GFP. E-H) Schematic strategy representation for patching selectively EP (E) and VP (H) neurons projecting to the LHb. Optic fiber placed above the STN and at the axon terminals, respectively. F-I) Example traces of STN-EP/VP responses to 1 ms optical stimulation every 30 s. Bath perfusion of AMPA glutamatergic receptor antagonists (D- AP5+DNQX) abolish excitary responses from the STN. G-J) Pie chart illustrarting the percentage of EP (G) and VP (J) cells responding to STN inputs. K) Population graph showing the amplitude of the excitatory postsynaptic currents (EPSCs) between EP (n=14) and VP (n=6) cells. Higher amplitudes on EP cells are reported (P=0.0010; MW- U test). H) Box plots for the latency of the EPSCs between EP and VP cells. Note smaller rates for EP cells (P=0.0031; MW-U test).

### Selective stimulation of STN terminals within VP induces place avoidance

Having identified connectivity between STN and VP, leading to activation in the LHb, we next investigated if photostimulation of STN-terminals in VP is sufficient to drive the aversive avoidance behavior observed upon direct STN stimulation. A new group of Pitx2^Cre^ mice were injected into the STN with the same AAV as previous mice, but this time, the fiber optic probes were bilaterally implanted above the VP instead of above the STN. These mice are referred to as STN^Pitx2^-VP/ChR2 (STN-VP mice); controls are STN^Pitx2^-VP/CTRL (Fig. 7). The response to STN-VP-photostimulation was tested using the same behavioral paradigms of affective behavior as implemented for STN-photostimulation (Opto-Open Field-induced grooming, Opto-Avoidance-EPM test, Opto-RT-CPP, Sugar-PR paradigm). This terminal-stimulation approach showed that selective photostimulation of STN terminals within VP caused a similar increase in face grooming as when STN cell bodies were stimulated (Suppl. Fig. 7). In the Opto-Avoidance-EPM test, STN^Pitx2^-VP/ChR2 mice showed decreased time spent and number of entries in the photostimulation-paired EPM arms as shown by STN mice (Fig. 7A and Suppl. Fig. 7).

**Figure 7.**
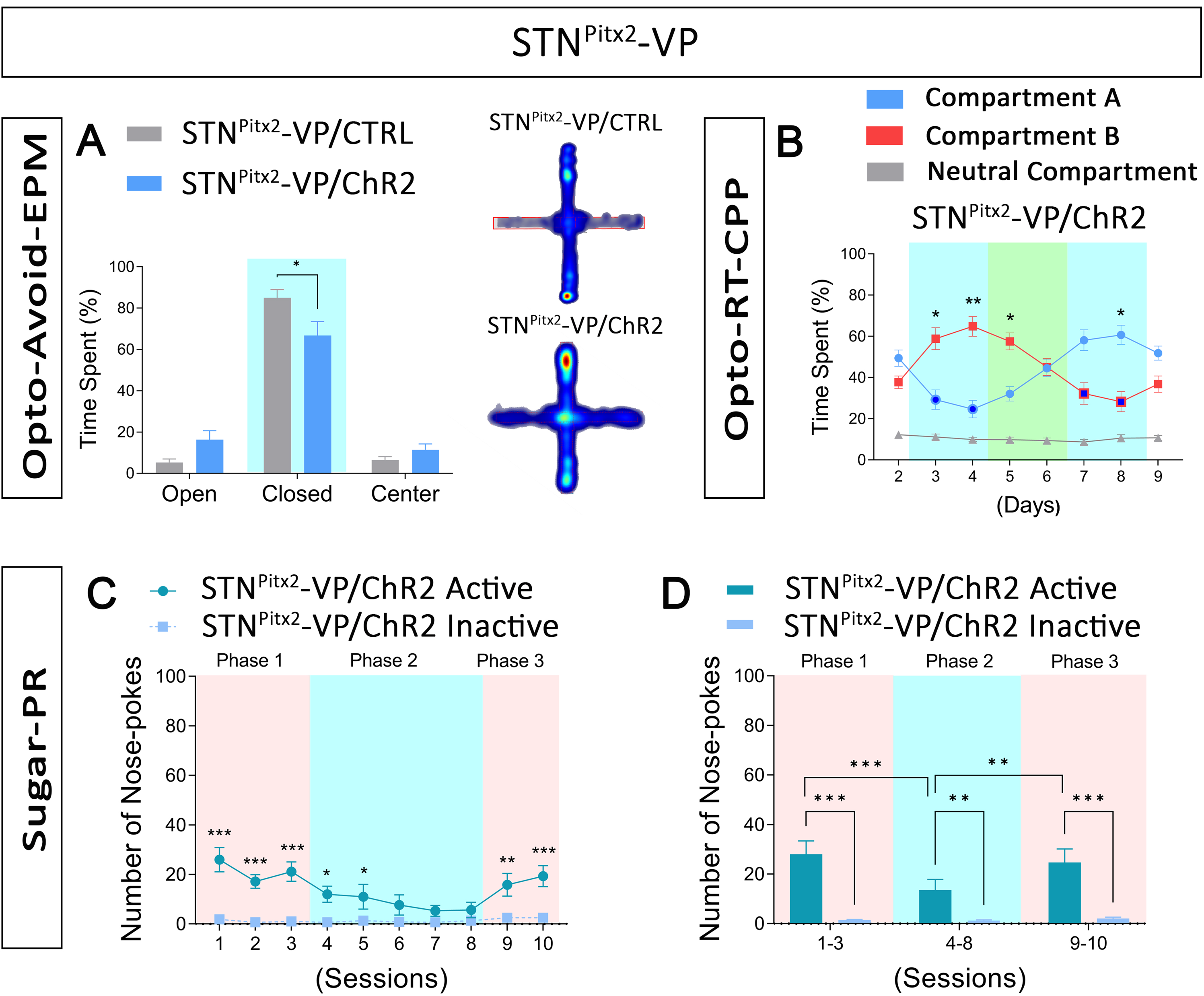
Selective photoactivation of STN-derived terminals within ventral pallidum (VP) is sufficient to drive place avoidance. (A-D) STN^Pitx2^ mice injected in STN as above, but fiber optic probes now implanted in VP instead of STN. Same battery of behavior tests as detailed above (Fig. 2, 3). A) Opto-Avoidance-EPM test. Time spent in arms: STN^Pitx2^-VP/CTRL (n=7) *vs.* STN^Pitx2^- VP/ChR2 (n=14), *p<0.05. Representative heat map showing occupancy for STN^Pitx2^- VP/CTRL and STN^Pitx2^-VP/ChR2. B) Opto-RT-CPP test. Percentage of time spent in each compartment: STN^Pitx2^-VP/ChR2 (n=14) compartment *A vs.* compartment B, *p<0.05, **p<0.01. Dark blue circles (Compartment A) and squares (Compartment B) indicate photostimulation (Light-ON) in that compartment. C) Number of STN^Pitx2^- VP/ChR2 (n=6) active *vs.* inactive nose-pokes: *p<0.05, **p<0.01, ***p<0.001. D) Average number of STN^Pitx2^-VP/ChR2 (n=6) active and inactive nose-pokes/phase: inactive *vs.* active, **p<0.01, ***p<0.001; STN^Pitx2^-VP/ChR2 active phase 1 *vs.* phase 2, ***p<0.001; STN^Pitx2^-VP/ChR2 active phase 2 vs. phase 3, **p<0.01. Suppl. Fig. 7 supporting data. (n) refers to number of mice. Data expressed as mean ±SEM.

In the Opto-RT-CPP paradigm (Fig. 7B), an active avoidance behavior to STN-VP- photostimulation was observed, with STN^Pitx2^-VP/ChR2 mice visiting and spending significantly less time in the light-paired compartment (A) than the light-unpaired compartment (B) both during the two days of real-time exposure and during the first conditioned test day (Fig. 7B, Suppl. Fig. 7). Upon reversal of light-pairing (STN^Pitx2^- VP/ChR2) mice reversed their preference by visiting and spending significantly more time in the now light-unpaired compartment (A) (Fig. 7B, Suppl. Fig. 7). Control mice showed an absence of photostimulation-dependent response throughout the sessions (Suppl. Fig. 7).

Finally, when STN-terminal stimulation was tested by placing STN^Pitx2^-VP/ChR2 mice in the sugar reinforcement paradigm (Fig. 7C, D), mice showed a similar response as STN-stimulated mice (STN^Pitx2^/ChR2 and STN^Vglut2^/ChR2) (Fig. 7C). However, the response was less pronounced. As STN-stimulated mice, also STN-VP-stimulated mice showed a significantly higher number of active compared to inactive NPs during the first phase and last phases with sugar only (Phase 1,3) (Fig. 7C, D). When exposed to STN-VP terminal photostimulation upon nose-poke for sugar, STN^Pitx2^-VP/ChR2 mice indeed showed avoidance, but their response was slower than that observed for STN-stimulated mice. This slower decrease in the number of active NPs led to maintained significance between active and inactive NPs when pooling all sessions in which mice received stimulation upon nose-poking for sugar (Fig. 7D). However, the number of active and inactive NPs reached similar values during the last 3 of 5 sessions (6-8) of Phase 2 (Fig. 7C) and the averaged number of active NPs during Phase 2 was significantly lower compared to both Phase 1 and Phase 3 (Fig. 7D). Taken together, the selective stimulation of STN-derived projection terminals within the VP verified the responses observed upon stimulation of the STN itself, thus identifying the STN-VP connection in behavioral avoidance.

## Discussion

The STN is a small thalamic brain structure described as critical to the central motor programs that enable us to move our body purposefully, allowing only relevant movement to be activated upon command. In this study, we identified aversion and avoidance-behavior as a direct consequence of optogenetic STN excitation in mice. This is an important finding considering the strategical position of the STN at the intersection between motor execution, and cognitive and affective function. Excitation of the immediately adjacent pSTN caused a different pattern of avoidance than STN excitation, in accordance with the different neurocircuitries identified for STN and pSTN, respectively. The findings of this study provide a neurobiological framework for emotional affect in the natural context and with implications for neurological and neuropsychiatric disorders in which STN dysfunction is a critical brain pathology.

STN is best known for, or at least most studied for, its role in movement inhibition in PD. However, PD contains both motor and non-motor symptom domains ^85–87^. PD is progressive in nature, and while no cure exists that restores movement control, treatments alleviating motor incapacity have existed for some decades. Of these, dopamine replacement therapy is the treatment of choice early in the disease, whereas DBS within, or near, the STN is a preferred treatment in late-stage PD ^9^. In contrast to the degeneration of dopamine neurons, there is no evidence for STN neuron degeneration in PD ^88^. Instead, the STN shows aberrant firing activity, which exacerbates the movement inability induced by loss of dopamine. STN-DBS improves PD motor symptoms, including brady- and akinesia, tremor, and rigidity ^89–91^. With STN-DBS, the effect is immediate: PD patients re-gain control over their body and can move around freely as long as the STN is stimulated. The effect is also reversible, demonstrating the crucial role of the STN in voluntary movement. Non-motor symptoms of PD include affective and cognitive symptoms that reduce the quality of life for PD patients beyond motor challenges. Non-motor symptoms have gained an interest over the past years, not least driven by the search for early biomarkers of PD, such as prodromal symptoms or altered brain activity that might allow earlier or more secure diagnosis, which in turn would motivate earlier treatment. In fact, anxiety and depression are affective non-motor symptoms of prodomal PD that may appear several years before motor symptoms allow diagnosis, and that today count as strong predictors of PD ^92–95^. Beyond PD, both STN-DBS and experimental STN-HFS have been shown to reduce compulsivity and drug consumption and relapse, leading to its use today as intervention of highly treatment-resistant OCD ^24, 96^, and to the proposed implementation of STN-DBS as treatment for substance use disorder ^35, 36^.

While STN-DBS can be highly beneficial, one major challenge is the emergence of motor, cognitive and affective side-effects ^10, 15, 97–100^. Such STN-DBS-induced side-effects include apathy and depression, neuropsychiatric symptoms that resemble affective symptoms of the non-motor symptom domain in PD. Of the reported side-effects, both men and women complain about depression upon STN-DBS ^101–104^. The neurobiological underpinnings correlating the STN with low mood state, and STN-DBS with depression, have remained elusive. This is partly due to poor information regarding the natural role of the STN even under healthy conditions, let alone during brain disorder in which STN dysfunction is implicated.

Aversion processing is essential to health and well-being, but its disturbed function can have deleterious effects for affected individuals. Aversive stimuli, such as the threat of a predator or the perception of an imminent dangerous situation induce the activation of a “survival state” designed to avoid or reduce the possible harmful outcome. In this context, aversive learning allows the animals to detect the aversive stimulus and learn to actively avoid it, while aversive Pavlovian conditioning serves to associate neutral stimuli to the hostile situation and environment reducing the probability of a related behavior being expressed ^105^. Obtaining a reward or avoiding punishment shapes decision-making and motivates learning with several brain regions playing a part is such vital responses ^105^. Patients with major depression disorder show mood-congruent biases in information-processing, implicating an association between depression and enhanced aversive pavlovian control over instrumental behavior ^106^.

In contrast to the recent interest in pallidal and habenular structures in aversion processing, the STN has been far less explored in this context. Main interest for the STN using the spatio-temporal specificity of optogenetics has focused around its role in motor control, given its key role in PD and additional movement disorders. Based on the strong interest in the therapeutic effects of STN-DBS, optogenetics in the STN has mainly been implemented in experimental animal models of PD ^107–115^. The natural role of STN in functions beyond induced PD-like symptoms has remained more poorly investigated. However, recent studies have explored the STN in locomotor control, action selection and stopping of cognitive processes ^57–59^.

Our current findings now identify aversive responses upon STN activation, possibly by acting as an upstream component of the LHb, a critical hub of aversion and mood regulation and clinically associated with depression. Indeed, STN excitation caused an evoked excitation of LHb neurons, demonstrating a common circuitry, but also found the association indirect. With the observed STN-mediated excitation of VP and EP neurons in our viral-genetic tracing combined with an electrophysiological strategy, these areas are here suggested to connect the STN and LHb. However, given the multitude of structures that communicate with the STN, additional experiments will be required to fully expose the STN circuitry of aversion. While further investigations will be needed to outline the neurocircuitry of STN-mediated aversion, the current findings clearly show that STN excitation caused both immediate place avoidance and cue-induced avoidance behavior, thus directly linking STN activation to aversion- processing. In this context, the STN, which in turn receives inputs from cortical regions supporting hedonic processing ^39, 116–118^, might play a modulatory role in decision-making in response to aversive conditions. Further, the current results pointed towards a different role for the adjacently located pSTN area in aversion processing; indeed immediate place avoidance was observed here which validated recent findings ^51^, but conditional aversion was not detected. Thus, any stimulation directed at pSTN would be expected to give a different affective outcome than STN stimulation. This finding further emphasizes the importance of neurocircuitry decoding of the STN and pSTN.

The STN projects directly to several of the structures recognized in aversive processing, including GPi/EP and VP ^2, 5, 58, 81–84, 119^, that in turn project to the LHb ^81–84^. Several of these different afferents to the LHb have proven responsible for mediating aversive behavior. For instance, optogenetic stimulation of glutamatergic neurons of the EP ^82, 83^, VP ^81, 84^ and LH ^52, 53^ projecting to the LHb induces place avoidance. In light of its important role integrating aversive and rewarding information ^62, 81, 120–124^, the VP might represent a plausible connecting structures to bridge STN and LHb in aversion processing. VP is mostly a GABAergic structure, but a recent study demonstrated the presence of glutamatergic neurons in the VP projecting to the LHb ^81, 84^. This glutamatergic population receives projections from the STN among other areas and selective stimulation of glutamatergic VP neurons evokes EPSCs in the LHb and induced a real-time avoidance ^84^. Now, our findings identifying optogenetically induced aversion upon stimulation of either the STN itself or its terminals in the VP, place the STN as a distinct brain nucleus engaged in aversion circuitry. Using *in vivo* extracellular recordings, we confirmed the activational response of the STN neurons themselves upon photostimulation, and also demonstrated that stimulation of STN glutamatergic neurons drive neuronal activity in the LHb, while patch-clamp results identify excitatory post-synaptic responses in EP and VP. Together, these electrophysiological findings provide strong support for the STN-VP-LHb pathway. However, with substantial projections between the STN and additional brain areas, including dopaminergic neurons of substantia nigra and ventral tegmental area that also have been associated with different kinds of aversion ^125, 126^, work remains to identity the complete neurocircuitry of the herein identified STN-mediated aversion.

The clinical importance of the STN highlights the need for decoding the role of this small nucleus in behavioral regulation. The STN is located in a narrow brain area, directly bordering the pSTN, recently implicated in both appetitive and aversive responses ^45–51^. The pSTN is of hypothalamic identity and shows vastly different projection pattern than the thalamic structure STN, as shown here and also in recent work from Kim et al ^47^. The pSTN^Tac1^ neuronal population projects heavily to limbic brain areas, including central amygdala, BNST, septum, while the STN^Pitx2^ shows far less projections in these areas and instead targets VP, EP, GP, SNr primarily. In the natural context, the STN and pSTN are distinct structures anatomically and functionally. However, one might consider that stimulating electrodes placed in the STN could affect immediately surrounding structures such as the pSTN, to which no specific anatomical border is present to guide stereotaxic surgery. With recent advances in transcriptomics, we recently identified *Tac1* as a selective promoter for pSTN ^65^, a finding which has been successfully implemented in recent study of the small and elusive pSTN ^45, 47, 50^. Here, taking advantage of a range of selective promoters to dissociate the STN and pSTN, we reasoned that a comparison between these anatomically associated structures would give clues to their roles in affective function, and hence provide information regarding their potential impact on DBS-induced side-effects. Indeed, by comparing projectivity and optogenetic excitation of STN and pSTN, their similar but yet clearly distinct roles in aversion and avoidance behavior was evident.

Further, not only adjacent structures, but also the functional heterogeneity of the STN itself poses challenges when electrodes are placed in the area and allowed to take control over the multitude of activities that are regulated here. Here, we used *Vglut2* and *Pitx2* promoters to direct opsin selectivity to the whole STN, and also tested *PV*, identified as promoter active in a subpopulation of STN neurons in both rodents and primates ^65, 67–70^. Based on the finding that STN excitation causes avoidance behavior, we speculated that the PV-expressing STN subpopulation might contribute to aversion-related function. Curiously, the STN^PV^ subpopulation mimicked some, but not all, behaviors assessed and was not concluded as a major player in behavioral avoidance. More work is needed to reveal the exact molecular identity of STN neurons engaging in the newly identified avoidance-type behavior which mice display in the current analyses.

To advance precision in treatment and reach symptom-alleviation without causing adverse side-effects, experimental strategies that enable revelation of the full repertoire of behaviors mediated by the subthalamic area are necessary. Considering that numerous adverse side-effects of STN-DBS that have been reported, including low mood state, depression, personality changes and even suicide ^13, 127^, manipulating the STN might come with a certain risk. While this issue has been challenging to resolve, any direct causality between STN excitation and behavioral aversion in mice is clearly crucial to take into consideration as putatively important not only experimentally, but also clinically. The present results underscore the importance of decoding the full repertoire of behavioral regulation executed by the STN, not least to improve treatment prediction and outcome in any intervention aiming to manipulate the STN and its pathways. As discussed above, STN-DBS has recently gained attention as treatment for additional neuropsychiatric disorders, including addiction ^35^. This interest is strongly based on findings that STN-DBS reduces symptoms of the dopamine dysregulation syndrome in PD ^128–131^. Also, studies in rodents showing that experimental STN-DBS induces different motivational responses depending on the nature of the reinforcers ^36, 64, 132–134^ and can reduce addictive behaviors ^36, 37, 135^, have identified a role for the STN which is relevant in addiction. The main obstacle to implementing this technique for addiction treatment seems to lie in the difficulty of recruiting human candidates ^136^. Further, low and high-frequency stimulation of the STN has different outcome in models of addiction ^32, 137^. However, it has also been shown that STN lesioning in rats leads to altered emotional state in response to various rewarding and aversive stimuli ^138^. Taken together, both experimental and clinical data clearly highlight the STN structure as critical in affective processing.

Here, we describe the identification of the STN as a non-canonical source of negative emotional value. Well embedded within the brain circuitry of negative reinforcing properties, the presented results point towards a pivotal role of the clinically relevant STN in aversive learning. Evidently, aberrant activity of STN circuitry, as well as its manipulation, may cause both beneficial and detrimental effects. Further studies will be needed to fully reveal the complete neurocircuitry engaging the STN in aversion processing.

## Supplemental file 1

Detailed description of statistics: Behavioral experiments.

## Acknowledgements

The authors thank James Martin, Baylor College of Medicine, Houston, Texas, USA, for generously providing the Pitx2^Cre^ transgenic mouse line, and Ole Kiehn, University of Copenhagen, Copenhagen, Denmark for generously providing the Vglut2^Cre^ transgenic mouse line. Marie Englund is thanked for technical assistance. This work was supported by Uppsala University and by grants to Å.W.M from the Swedish Research Council (Vetenskapsrådet), the Swedish Brain Foundation (Hjärnfonden), Parkinsonfonden, and the Research Foundations of Bertil Hållsten, OE & Edla Johansson, Zoologiska and Åhlén. This work has benefited from a government grant awarded to the University of Bordeaux as an Initiative of Excellence, under the France 2030 plan (J.B and Å.W.M).

## Supplemental file 1

Detailed description of the statistics used for the different behavioral experiments.

---

**Supplemental Figure 1.**
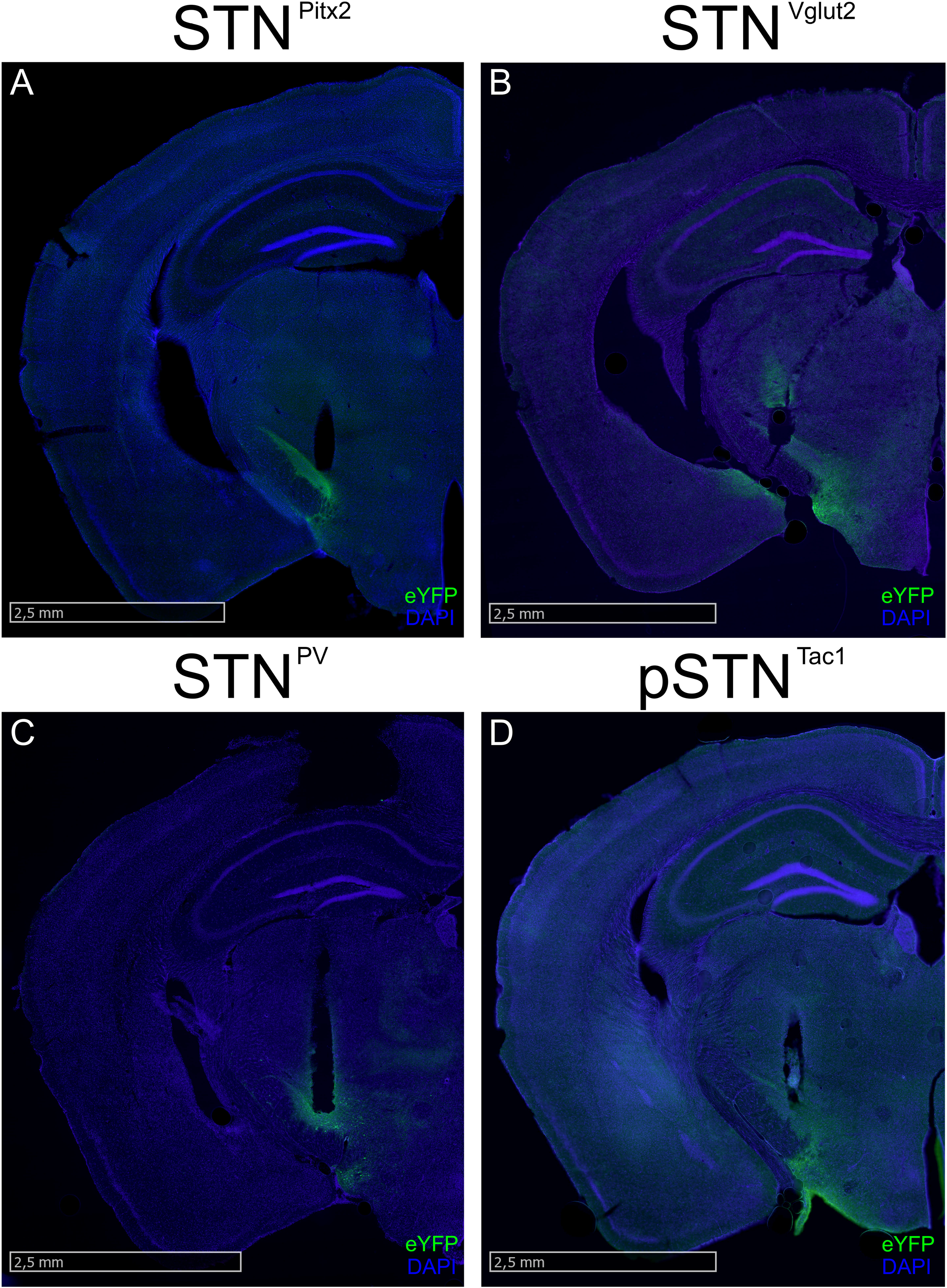
Representative image of coronal sections showing YFP fluorescence (green) in cell bodies of: A) STN^Pitx2^/CTRL, B) STN^Vglut2^/CTRL, C) STN^PV^/CTRL and D) pSTN^Tac1^/CTRL mice with optic cannula track above the STN; nuclear marker (DAPI, blue); scale, 2,5 mm.

**Supplemental Figure 2.**
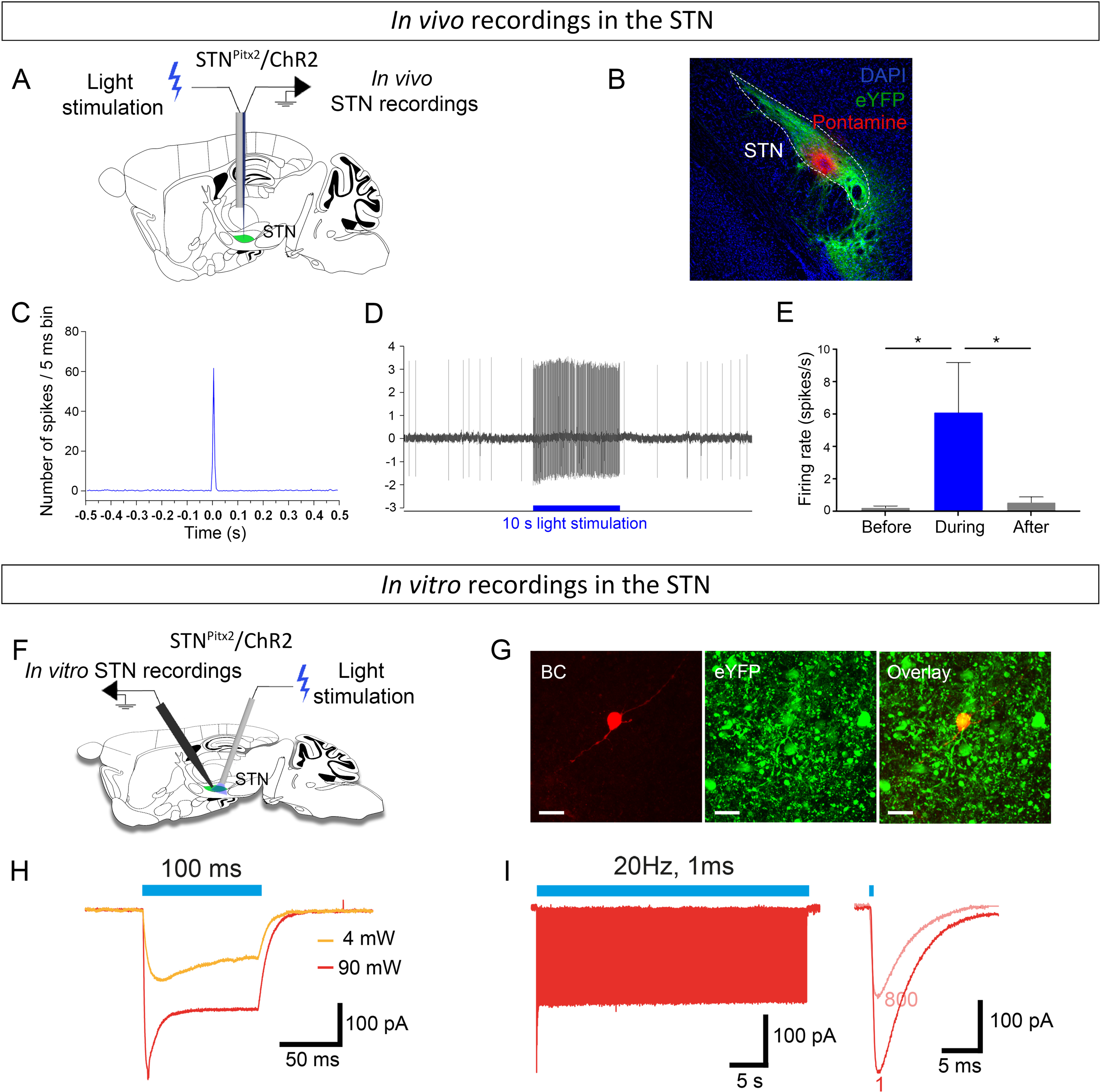
A) Procedure for STN optotagging. B) YFP-positive neurons (green) in the STN; pontamine blue sky deposit (red dot) at the last recorded coordinate; nuclear marker (DAPI, blue) (scale, 200 µm). C) Averaged PSTH of STN-excited neurons upon light stimulation (0.5 Hz, 5 ms pulse duration, 5-8 mW). D) Example of an excited STN neuron during a behavioral protocol (20 Hz, 5 ms pulse duration, 5-8 mW) with the frequency and the recording trace. E) Frequency of STN neurons before, during and after 10 s light stimulation with behavioral protocol (20 Hz, 5 ms pulse duration, 5-8 mW); data expressed as mean ± SEM, n=7, both *p=0.15. F) Representation of para- sagittal slice with patch-clamp electrode and optic fiber placed above the STN. G) Confocal images of STN neuron filled with biocytin (BC) and expressing the ChR2- YFP construct (visualized by YFP). Scale bar: 10µm. H) Representative example of typical ChR2-mediated currents induced by a 1 s blue light (λ = 470 nm) pulses at intensities of 4mW and 90 mW, respectively. I) ChR2-mediated currents induced by a train of blue light of 40 s duration at 20 Hz (individual light pulses of 1 ms). Abbreviations: STN, subthalamic nucleus; VP, ventral pallidum; GP, globus pallidus, EP; entopenduncular nucleus; SNr, substantia nigra pars reticulata; SNc, substantia nigra pars compacta.

**Supplemental Figure 3.**
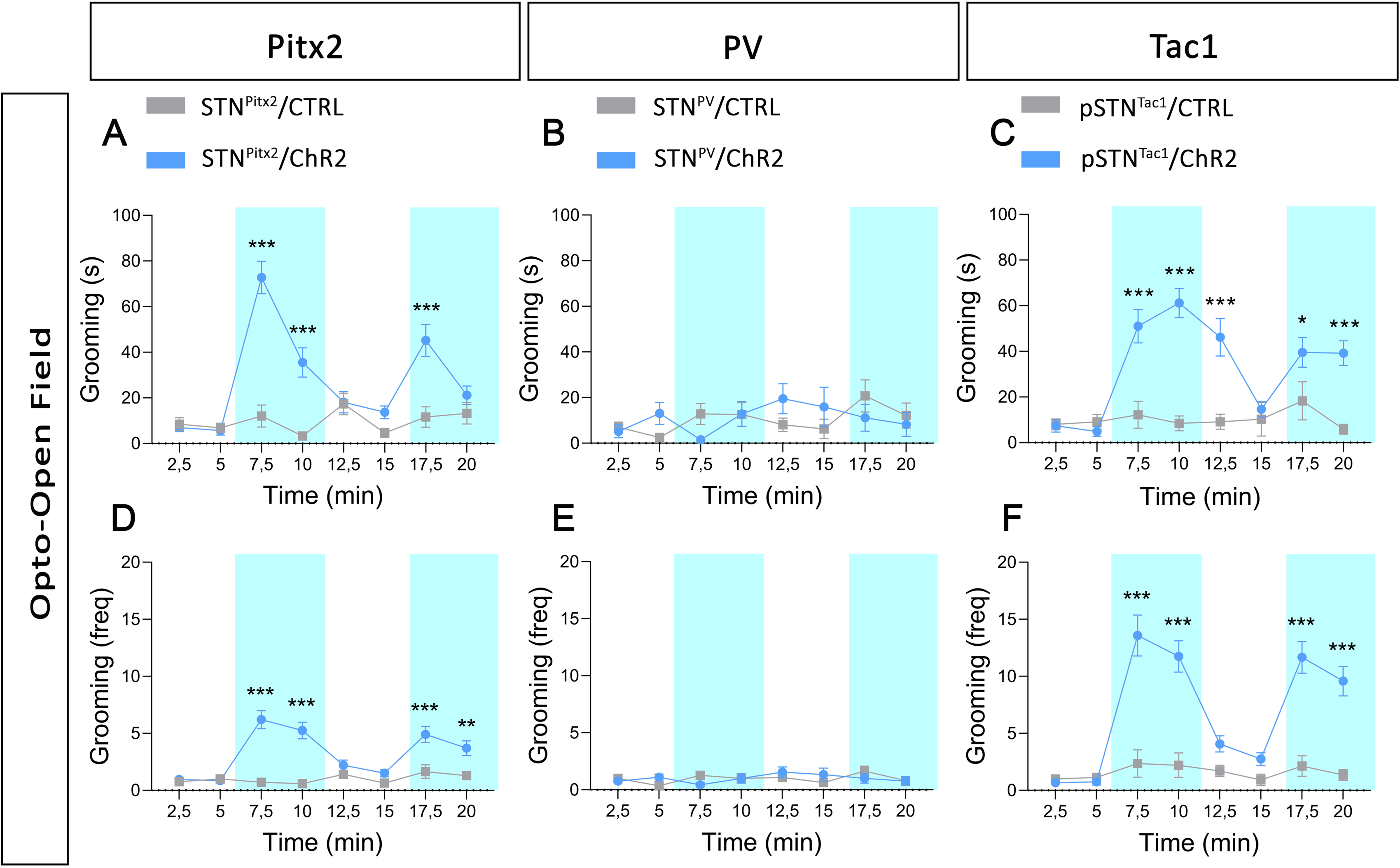
Time spent grooming: CTRL vs. ChR2 *p<0.05, **p<0.01, ***p<0.001, divided in 2.5 min intervals for: A) STN^Pitx2^/ChR2 (n=20) and STN^Pitx2^/CTRL (n=17) mice; B) STN^PV^/ChR2 (n=9) and STN^PV^/CTRL (n=11) mice; C) pSTN^Tac1^/ChR2 (n=12) and pSTN^Tac1^/CTRL (n=14) mice. Frequency of grooming: CTRL vs. ChR2 **p<0.01, ***p<0.001, divided in 2.5 min intervals for: D) STN^Pitx2^/ChR2 (n=20) and STN^Pitx2^/CTRL (n=17) mice; E) STN^PV^/ChR2 (n=9) and STN^PV^/CTRL (n=11) mice; F) pSTN^Tac1^/ChR2 (n=12) and pSTN^Tac1^/CTRL (n=14) mice. Data expressed as mean ±SEM.

**Supplemental Figure 4.**
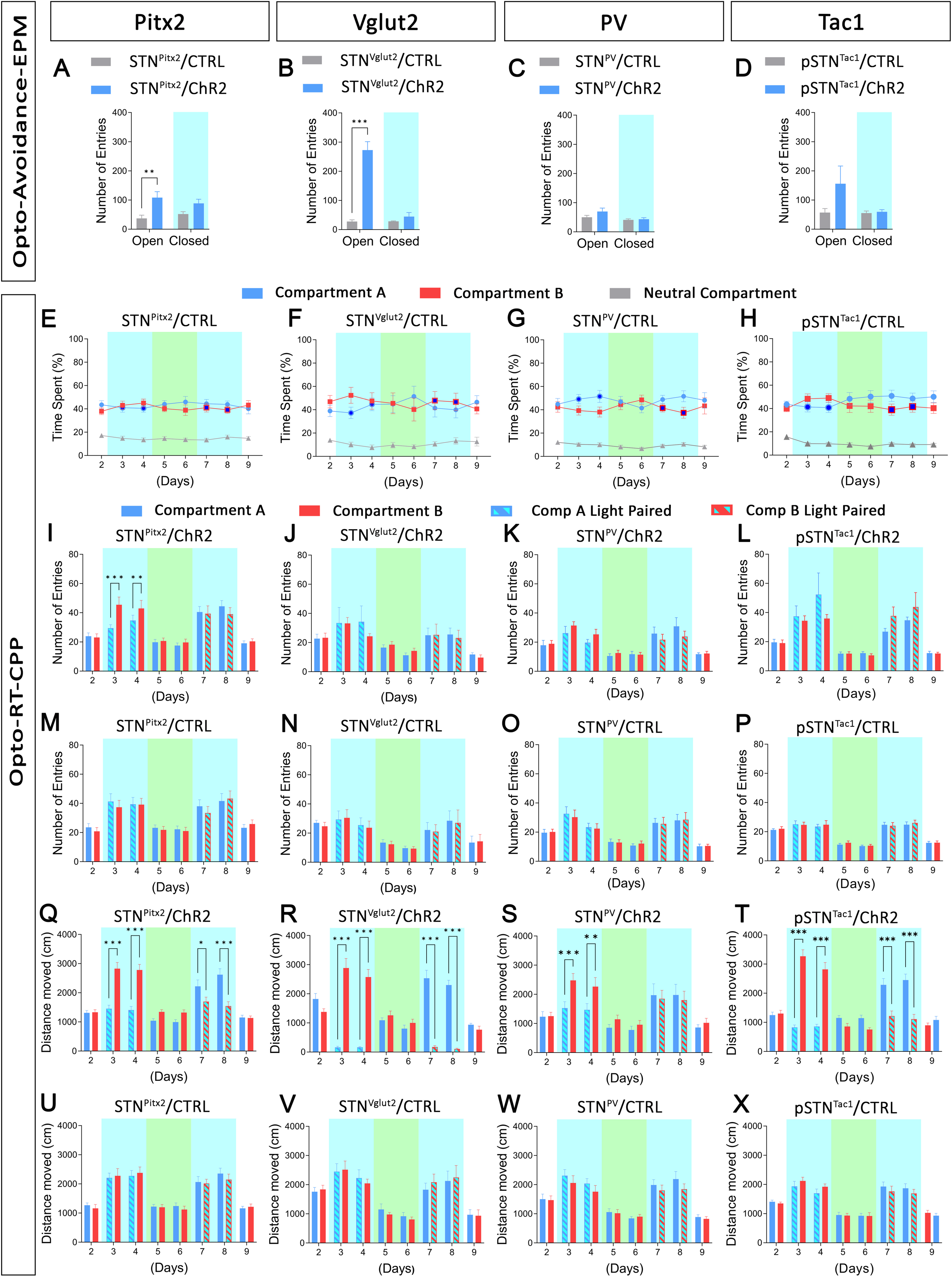
Opto-Avoidance-EPM test, number of entries in arms: open arms CTRL vs. ChR2, **p<0.01, ***p<0.001: A) STN^Pitx2^/ChR2 (n=15) and STN^Pitx2^/CTRL (n=14) control mice; B) STN^Vglut2^/ChR2 (n=7) and STN^Vglut2^/CTRL (n=6); C) STN^PV^/ChR2 (n=9) and STN^PV^/CTRL (n=11) control mice; D) pSTN^Tac1^/ChR2 (n=11) and pSTN^Tac1^/CTRL (n=12) control mice. Opto-RT-CPP, percentage of time that mice spent in each compartment: compartment A vs. compartment B. Dark blue circles (compartment A) and squares (compartment B) indicate photostimulation in that compartment: E) STN^Pitx2^/CTRL (n=13) mice; F) STN^Vglut2^/CTRL mice (n=6); G) STN^PV^/CTRL mice (n=11); H) pSTN^Tac1^/CTRL (n=12) mice. Opto-RT-CPP, number of entries in each compartment: compartment A vs. compartment B, **p<0.01, ***p<0.001: I) STN^Pitx2^/ChR2 (n=15) mice; J) STN^Vglut2^/ChR2 (n=7) mice; K) STN^PV^/ChR2 (n=9) mice; L) pSTN^Tac1^/ChR2 (n=11) mice; M) STN^Pitx2^/CTRL (n=13) mice; N) STN^Vglut2^/CTRL (n=6) mice; O) STN^PV^/CTRL (n=11) mice; P) pSTN^Tac1^/CTRL (n=12) mice. Distance moved in each compartment: compartment A vs. compartment B, *p<0.05, **p<0.01, ***p<0.001: Q) STN^Pitx2^/ChR2 (n=15) mice; R) STN^Vglut2^/ChR2 (n=7) mice; S) STN^PV^/ChR2 (n=9) mice; T) pSTN^Tac1^/ChR2 (n=11) mice; U) STN^Pitx2^/CTRL (n=13) mice; V) STN^Vglut2^/CTRL (n=6) mice; W) STN^PV^/CTRL (n=11) mice; X) pSTN^Tac1^/CTRL (n=12) mice. Data expressed as mean ±SEM.

**Supplemental Figure 5.**
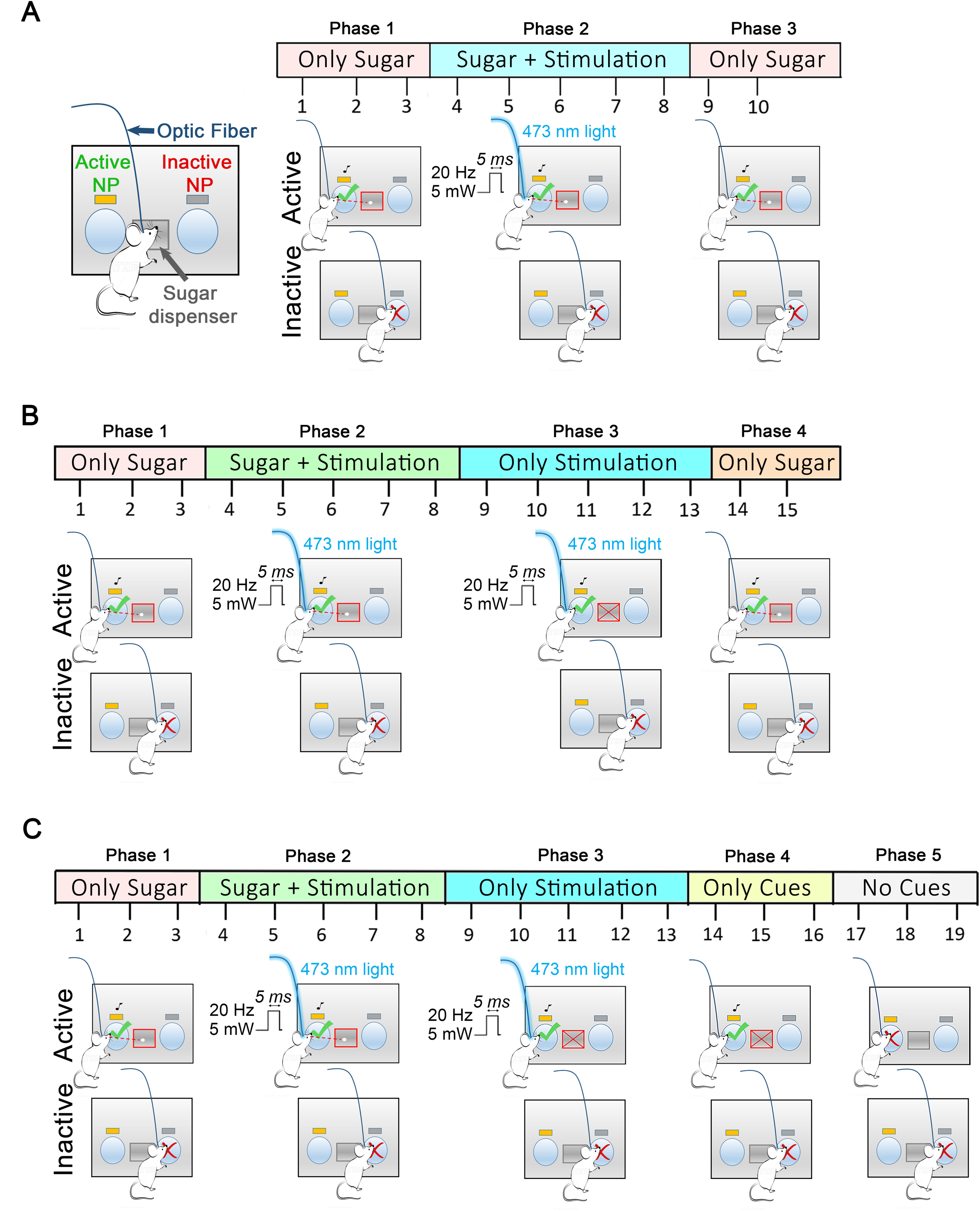
Graphical representations of Sugar-PR protocols used for: A) STN^Pitx2^ and STN^Vglut2^ mice; B) STN^PV^ mice; C) pSTN^Tac1^ mice.

**Supplemental Figure 6.**
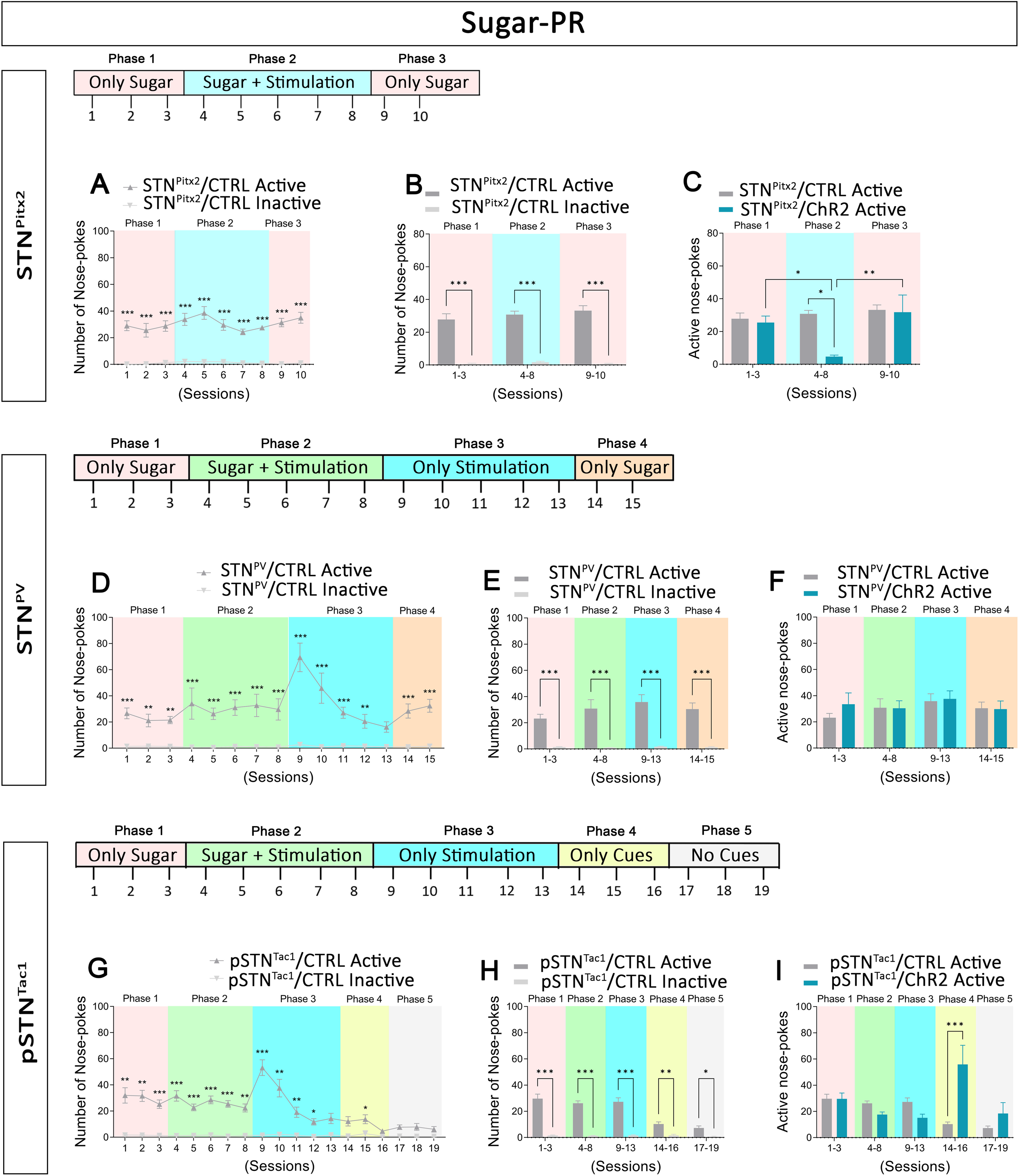
A) Number of STN^Pitx2^/CTRL (n=5) active vs. inactive nose-pokes: ***p<0.001. B) Average number of STN^Pitx2^/CTRL (n=5) active and inactive nose-pokes/phase: phase 1 inactive vs. active, ***p<0.001; phase 2 inactive vs. active ***p<0.001; phase 3 inactive vs. active ***p<0.001. C) Average number of active nose-pokes/phase: phase 2 STN^Pitx2^/CTRL (n=5) vs. STN^Pitx2^/ChR2 (n=8), *p<0.05; STN^Pitx2^/ChR2 phase 1 vs. phase 2, *p<0.05; STN^Pitx2^/ChR2 phase 2 vs. phase 3, **p<0.01. D) Number of STN^PV^/CTRL (n=9) active vs. inactive nose-pokes: **p<0.01, ***p<0.001. E) Average number of STN^PV^/CTRL (n=9) active and inactive nose-pokes/phase: inactive vs. active, ***p<0.001. F) Average number of STN^PV^/CTRL (n=9) vs. STN^PV^/ChR2 (n=9) active nose-pokes/phase. G) Number of pSTN^Tac1^/CTRL (n=10) active vs inactive nose-pokes: *p<0.05, **p<0.01, ***p<0.001. H) Average number of pSTN^Tac1^/CTRL (n=10) active and inactive nose-pokes/phase: inactive vs. active, *p<0.05, **p<0.01, ***p<0.001. I) Average number of pSTN^Tac1^/CTRL (n=10) vs. pSTN^Tac1^/ChR2 (n=11) active nose-pokes/phase: ***p<0.001. Data expressed as mean ±SEM.

**Supplemental Figure 7.**
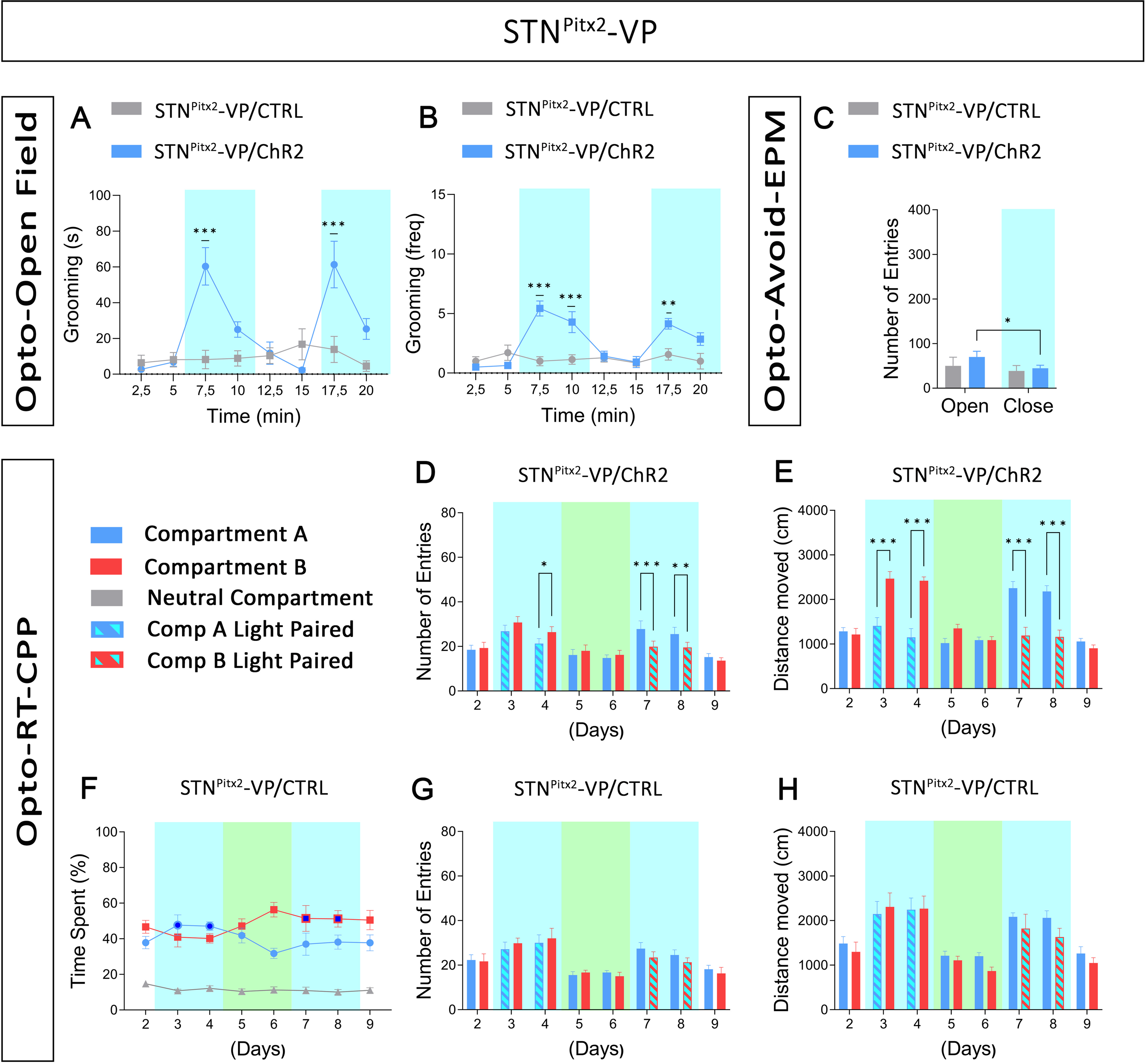
A) Time spent grooming: STN^Pitx2^-VP/CTRL (n=7) vs. STN^Pitx2^-VP/ChR2 (n=14) ***p<0.001, divided in 2.5 min intervals B) Frequency of grooming: STN^Pitx2^-VP/CTRL (n=7) vs. STN^Pitx2^-VP/ChR2 (n=14) **p<0.01, ***p<0.001. C) Opto-Avoidance-EPM, number of entries in arms STN^Pitx2^-VP/ChR2 (n=14) vs. STN^Pitx2^-VP/CTRL (n=7): STN^Pitx2^-VP/ChR2 open vs. closed arms, *p<0.05. D) Opto-RT-CPP, number of entries in each compartment: STN^Pitx2^-VP/ChR2 (n=14) compartment A vs. compartment B, *p<0.05, **p<0.01, ***p<0.001. E) Opto-RT-CPP, distance moved in each compartment: STN^Pitx2^-VP/ChR2 (n=14) compartment A vs. compartment B, ***p<0.001. F) Opto-RT-CPP, percentage of time that mice spent in each compartment: STN^Pitx2^-VP/CTRL (n=7). Dark blue filled circles (compartment A) and squares (compartment B) indicate photostimulation in that compartment. G) Opto-RT- CPP, number of entries in each compartment: STN^Pitx2^-VP/CTRL (n=7). H) Opto-RT- CPP, distance moved in each compartment: STN^Pitx2^-VP/CTRL (n=7). Data expressed as mean ±SEM.

## METHODS SECTION

### Mice: Source, housing and ethics

Sources of Cre-recombinase mouse strains: Vglut2^Cre^ ^1^ (courtesy of Ole Kiehn, Copenhagen University, Denmark), Pitx2^Cre^ ^2, 3^ (courtesy of James Martin, University of Texas, USA), Tac1^Cre^ ^4^ (Jackson Laboratories, B6;129S-Tac1^tm1.1^(cre)^Hze^/J; Jax #021877), PV^Cre 5^ (Jackson Laboratories, B6;129P2-Pvalb^tm1^(cre)^Arbr^/J; Jax #017320, common name B6 PV^Cre^). Adult mice of both male and female sex were analysed.

Vglut2^Cre^, Pitx2^Cre^, Tac1^Cre^, PV^Cre^ transgenic mice were maintained at the local animal facility of Uppsala University before and during behavioral experiments, or at University of Bordeaux (Pitx2^Cre^, Vglut2^Cre^) for *in vivo* electrophysiological experiments. Mice were maintained on a C57BL/6NTac background, and all breedings were performed *in-house*. Mice had access to food and water *ad libitum* in standard humidity and temperature conditions and with a 12-hour dark/light cycle. PCR analyses were performed on ear biopsies to confirm the Cre-positive genotype (Vglut2^Cre^ forward; 5’TTGCATCGCATTGTCTGAGTAG, reverse; 5’TTCCCACACAAGATACAGACTCC, Pitx2^Cre^, Tac1^Cre^, PV^Cre^ forward, 5’GCG GTC TGG CAG TAA AAA CTA TC; reverse, 5’GTG AAA CAG CAT TGC TGT CAC TT). All experimental procedures using mice followed Swedish (Animal Welfare Act SFS 1998:56) and European Union Legislation (Convention ETS 123 and Directive 2010/63/EU), and were approved by local ethical committees in Uppsala and/or Bordeaux (N°50120205-A).

### *In situ* hybridization histochemistry Brain preparation

To prepare brain sections for mRNA analysis by fluorescent *in situ* hybridization (FISH), brains were quickly dissected from mice euthanized by cervical dislocation, snap-frozen in cold (-30°C to -35°C) 2-methylbutane (≥99%, Honeywell) and kept at - 80°C until usage. Coronal serial sections were prepared on a cryostat at 16µm thickness and placed onto Superfrost glass slides in series of 8 slides (Thermo Fischer). Slides prepared for FISH analysis were stored in -80°C until usage.

### Fluorescent *in situ* hybridization (FISH) analysis for detection of mRNA in brain sections

For detection of selected mRNAs in cryosections, mRNA-directed fluorescent riboprobes were prepared from cDNA (Oramacell, Paris). Note: For FISH analyses, mice at postnatal day (P) 28 were used. Probes were synthesized by PCR from DNA template containing T3 and T7 promotor sequences using promoter-attached primers to detect: Vglut2: NM_080853.3; bases 2315-3244; Pitx2: NM_001042504.2; bases 792-1579 Tac1: NM_009311.3; bases 36-77, PV: NM_001330686.1; bases 92-591. Digoxigenin (DIG) and fluorescein-labeled RNA probes were made by a transcriptional reaction with incorporation of DIG or fluorescein labeled nucleotides^6^. Specificity of probes was verified using NCBI blast.

FISH experiments were performed following our previously described protocol ^6^. Cryosections were air-dried, fixed in 4% paraformaldehyde and acetylated in 0.25% acetic anhydride/100 mM triethanolamine (pH 8). Sections were hybridized for 18 h at 65°C in 100 µl of formamide-buffer containing 1 µg/ml digoxigenin-labeled riboprobe (DIG) and 1 µg/ml fluorescein-labeled riboprobe. Sections were washed at 65°C with SSC buffers of decreasing strength, and blocked with 20% fetal bovine serum and 1% blocking solution. Fluorescein epitopes were detected with horseradish peroxidase (HRP) conjugated anti-fluorescein antibody at 1:5000 and revealed using Cy2- tyramide at 1:250. HRP-activity was stopped by incubation of sections in 0.1 M glycine followed by a 3% H2O2 treatment. DIG epitopes were detected with HRP anti-DIG Fab fragments at 1:1500 and revealed using Cy3 tyramide at 1:100. Nuclear staining was performed with 4’ 6-diamidino-2-phenylindole (DAPI. All slides were scanned at 20x resolution using the NanoZoomer 2.0-HT (Hamamatsu, Japan). Laser intensity and time of acquisition were set separately for each riboprobe. Images were analyzed using the NDP.view2 software (Hamamatsu Photonics). Regions of interest were identified according to the Paxinos mouse brain atlas ^7^. Positive cells refer to a staining in a cell body clearly above background and surrounding a DAPI-stained nucleus. Colocalization was determined by the presence of the signals for both probes in the soma of the same cell.

### Stereotaxic virus injection and fiber optic probe implantation

*Virus injection*: Stereotaxic injections were performed in Pitx2^Cre^, Vglut2^Cre^, PV^Cre^, Tac1^Cre^ mice under isofluorane anesthesia (4% for induction and maintained at 1-1.5% air mix v/v). After being placed in a stereotaxic apparatus, mice received sub- cutaneous injection of analgesic and anti-inflammatory drugs (buprenorphine, 0.1 mg/Kg and Carprofen, 5 mg/Kg) as well as a local analgesic (lidocaine, 7 mg/Kg) before the incision of the skin. Mice were bilaterally injected in the STN/pSTN with a virus containing either a Cre-dependent Channelrhodopsin (ChR2) construct coupled with an eYFP reporter (rAAV2/EF1a-DIO-hChR2(H134R)-eYFP), or only the eYFP reporter (rAAV2/EF1a-DIO-eYFP), respectively, 3.8x10^12^ virus molecules/mL and 4.6x10^12^ virus molecules/mL (viruses purchased from UNC Vector Core, Chapel Hill, NC, USA) at the following mouse brain coordinates (from Paxinos and Franklin, 2013): anteroposterior (AP) = -1.90 mm, mediolateral (ML) = +/- 1.70 mm from the midline and 250 nL of virus was injected with a NanoFil syringe (World Precision Instruments, Sarasota, FL, USA) at two dorsoventral levels (DV) = -4.65 mm and -4.25 mm from the dura matter at 100 nL.min-1. *Optic cannula implantation*: Optic cannulas (Doric Lenses) were implanted directly after completion of virus injections in mice preparated for behavior analysis. Two skull screws were implanted in the skull to hold the optic cannula-cement-skull complex. Primers were then applied and harden with UV light (Optibond). Optic cannulas were implanted bilaterally above the STN (coordinates: AP = -1.90 mm, ML = +/-1.70 mm from the midline DV = -4.30 mm) or the VP (coordinates: AP = +0.45 mm, ML = +/-1.55 mm from the midline, DV = -4.00 mm), and fixed with dental cement. 1 mL of saline was injected subcutaneously at the end of the surgery.

For *ex vivo* electrophysiology experiments, Vglut2^Cre^ mice were injected into the LHb and the STN. A glass pipette (30-50 µm tip diameter) filled with a rgAAV2-hSyn-DIO-EGFP and was lowered at the LHb coordinates: AP: -1.55 mm; ML: -0.50 mm; DV: -2.65 mm depth and 120 nL were injected using a picospritzer III (Parker). Similarly, a glass pipette filled with AAV2-EF1a-DIO-ChR2(H134R)-mCherry was lowered at the STN coordinates: AP: -1.90 mm; ML: -1.60 mm; DV: -4.65/-4.30 mm and 2 times 120 nL were injected. 5 min after the last injection, the glass pipette was withdrawn. After stitching, mice were transferred to a cage placed on a heating pad until waking up.

### Single-cell extracellular recordings

*Surgery*: *In vivo* single cell extracellular recordings started after a post-injection recovery period of at least 4 weeks. Mice were anesthetized with a mix isoflurane-air (0.8-1.8 % v/v) and placed in a stereotaxic apparatus. Optic fiber, optrode and glass micropipettes coordinates were AP = -1.90 mm, ML = +/- 1.70 mm and DV = -4.30 mm for the STN and AP = -1.60 mm, ML = +/-0.50 mm, DV = -2.00 to -3.30 mm for the LHb.

*STN Optotagging:* A custom-made optrode was made with an optic fiber (100 µm diameter, Thorlabs) connected to a laser (MBL-III-473 nm-100 mW laser, CNI Lasers, Changchun, China) mounted and glued on the recording glass micropipette which was filled with 2% pontamine sky blue in 0.5 µM sodium acetate (tip diameter 1-2 µm, resistance 10-15 MΩ). The distance between the end of the optic fiber and the tip of the recording pipette varied between 650 nm and 950 nm. Extracellular action potentials were recorded and amplified with an Axoclamp-2B and filtered (300 Hz/0.5 kHz). Single extracellular spikes were collected online (CED 1401, SPIKE2; Cambridge Electronic Design). The laser power was measured before starting each experiment using a power meter (Thorlabs). The baseline was recorded for 100 seconds for each neuron before starting any light stimulation protocols which were set and triggered with Spike2 software. Light protocol consisted in a peristimulus time histogram (PSTH, 0.5 Hz, 5 ms pulse duration, 5-8 mW) for at least 100 seconds followed, after returned to baseline, by a “behavioral” protocol corresponding to the parameters used for behavioral experiments (20 Hz, 5 ms pulse duration, 5-8 mW).

*LHb recordings:* An optic fiber and a recording pipette filled with either 2% pontamine sky blue or 2% neurobiotin were respectively positioned in the STN and LHb. For each LHb neurons, a PSTH was recorded upon STN optogenetic stimulation (PSTH 0.5 Hz, 5 ms pulses, 5-8 mW) for at least 100 seconds. Neurons are considered as excited during the PSTH protocol when, following the light pulses centered on 0, the number of spikes/5ms bin is higher than the baseline (-500 ms to 0 ms) plus two times the standard deviation. Injection of neurobiotin (2% neurobiotin in 0.5 M acetate sodium) by juxtacellular iontophoresis was performed in some cases for precise spatial identification of excited neurons.

### *Ex vivo* electrophysiology

STN, EP and VP neurons were recorded in acute brain slices, prepared as previously described ^8, 9^. Briefly, STN^Pitx2^/CTRL, STN^Pitx2^/ChR2 and STN^Vglut2^ mice (> 12 months) were deeply anesthetized with an i.p. injection of ketamine/xylazine (75/10 mg/kg) mixture and then perfused trans-cardially with ice-cold (0-4°C) modified artificial cerebrospinal fluid (ACSF), equilibrated with 95% O_2_ and 5% CO_2_, and containing (in mM): 230 sucrose, 26 NaHCO_3_, 2.5 KCl, 1.25 NaH_2_PO_4_, 0.5 CaCl_2_, 10 MgSO_4_ and 10 glucose. (pH∼7,35). The brain was quickly removed, glued to the stage of a vibratome (VT1200S; Leica Microsystems, Germany), immersed in the ice-cold ACSF and sectioned into 300 μm thick parasagittal slices. Slices containing the LHb, the STN and the VP/EP were then incubated for 1 hour to a standard oxygenated ACSF solution, warmed (∼35°C) containing (in mM unless otherwise stated): 126 NaCl, 26 NaHCO_3_, 2.5 KCl, 1.25 NaH_2_PO_4_, 2 CaCl_2_, 2 MgSO_4_, 10 glucose, 1 sodium pyruvate and 4.9 μML-gluthathione reduced. Single slices were then transferred to a recording chamber at room temperature 22-26 °C and perfused continuously with oxygenated ACSF.

### Electrophysiological recordings

Patch-clamp recordings were performed under infrared gradient contrast video microscopy on an upright microscope (E600FN, Eclipse workstation, Nikon, Japan) equipped with a 60x water-immersion objective (Nikon Fluor 60 X/1.0 NA). The microscope was also equipped with epifluorescence (Nikon Intensilight C-HGFI) allowing visualization of STN neurons expressing the ChR2-mCherry as well as LHb-projecting EP and VP neurons retrogradely labelled with GFP, respectively.

STN, VP and EP neurons were recorded using low-resistance pipettes (impedance, 3–8 MΩ) prepared from borosilicate glass capillaries (GC150F10; Warner Instruments, Hamden, CT, USA) with a horizontal puller (P- 97; Sutter Instruments, Novato, CA, USA). In voltage-clamp experiments, the internal solution contained the following (in mM): 135 K-gluconate, 3.8 NaCl, 1 MgCl_2_.6H_2_O, 10 HEPES, 0.1 EGTA, 0.4 Na_2_GTP, 2 Mg_1.5_ATP, 5 QX-314 and 5.4 biocytin (pH=7.2, ∼292 mOsm). Data were acquired using a Multiclamp 700B amplifier connected to a Digidata 1550B digitizer controlled by Clampex 10.3 (Molecular Devices, Sunnyvale, CA, USA). Acquisitions were performed at 20 KHz and low-pass filtered at 4 KHz. Series resistance was monitored throughout the experiment by voltage steps of -5 mV. Data were discarded when the series resistance changed by >20%. Biocytin-filled neurons were identified.

For optogenetic stimulation, a LED laser source (Prizmatix, Israel) connected to optic fiber (∅: 500 µm) was placed above the brain slice. Light intensities ranged from 4mW to 90mW at the tip of the optic fiber depending of the type experiments. For cell body stimulation, continuous 100 ms long duration light stimulation was applied at low (4mW) and high (90 mW) intensities. To evoke synaptic transmission, single pulses or train of stimulation (800 pulses at 20Hz) of 1ms duration at full power (90 mW) were used in order to maximize axon terminal depolarization and efficient release of neurotransmitter. The magnitude of EPSCs was examined in whole-cell voltage-clamp mode at a holding potential of -60 mV. Junction potential (∼13 mV) was not corrected. EPSC latencies were calculated as the difference between the time of the pulse and the beginning of the negative deflection of the EPSC. After electrophysiological recordings, slices were fixed overnight in 4% paraformaldehyde, and maintained in PBS-azide at 0.2% at 4°C until immunohistochemical processing.

### Drugs

Unless otherwise stated, all pharmaceutical substances were prepared in distilled water as concentrated stock solutions and stored at -20°C. On the day of the experiment, the substances were diluted and applied through the bath perfusion system. In some experiments glutamatergic antagonists were used to demonstrate that the recorded EPSCs from STN inputs were mediated by AMPA/Kainate and MNDA receptors. NMDA receptors were blocked with 50 μM D-(–)-2-amino-5- phosphonopentanoic acid (D-APV). AMPA/kainate receptors with 20 μM 6,7- dinitroquinoxaline-2,3-dione (DNQX disodium salt).

### Behavioral testing

Following a post-surgical recovery period of approximately four weeks, mice were analyzed in a battery of behavioral tests. Throughout, the following stimulation protocol was used: 473 nm light, 5 mW, 20 Hz, 5 ms pulse delivered by a MBL-III-473 nm-100 mW laser (CNI Lasers, Changchun, China). Duration and condition of stimulation are specified for each test. After completed behavioral tests, mice were sacrificed and brains analysed histologically and for assessment of optic cannula position (see below for details). The initial sample size was determined based on previous published studies with similar experimental design.

### Criteria for exclusion from the analysis

Mice were excluded from individual test analysis if:

1. In the Opto-Open Field they jumped out of the arena during photostimulation so that the analysis of other parameters was made impossible.
2. In the Opto-RT-CPP if, during the pre-test (Day 1), there was a strong initial preference, such that the time spent in a compartment exceeded 80% of the total, or a strong avoidance, so that the time spent in a compartment was less than 20% of the total.
3. During Sugar-PR if they have not acquired a stable rate of self-sugaring, which consisted of at least 10 sugar lozenges received/session with <20% variation during three consecutive days or if they have done more than 30 active nose-pokes in a short time during the initial sessions of the paradigm.
4. In Opto-Avoidance-EPM if the mice were immobilized due to entanglement of the optical fibers during the test.

Mice were excluded from all analyzes if post-hoc histological examination revealed insufficient viral expression or the cannulas were not close enough to the site of interest. Mice were excluded if histological analysis was not possible due to premature death of the animal or if the animal had to be sacrificed for health reasons during the study. In case of loss of the optical assembly during the study, only the data acquired up to that moment were included. In one case the animal was excluded because an incorrect genotype was revealed during post-hoc analysis.

### Habituation

Three weeks after surgery and before the first behavioral test, all mice were handled and habituated to the experimental room and to the optic cables to reduce the stress during the day of the experiment. Before each behavioral test, mice were acclimatized for 30 minutes in the experimental room.

### Opto-Open Field

Mice were individually placed in neutral cages for 3 minutes in order to recover after connecting the optic cables. Mice were subsequently placed in the central zone of the open field arena and allowed to freely explore it for 5 min before starting the test. The open-field chamber consisted in a 50 cm, squared, transparent, plastic arena with a white floor. The Opto-Open Field test consisted of a 20-minute session divided into 5- minute alternating time intervals (OFF-ON-OFF-ON). Each 5-minute interval was then divided into two 2.5-minute intervals for analysis. The patch cable was connected to a rotary joint, which was attached on the other end to a laser that was controlled by an Arduino Uno card. During the light ON trials, blue light was delivered according to the behavioral stimulation protocol. Self-grooming and escape behaviors were manually recorded by an experimenter blind to the experimental groups using the EthoVision XT 14.0 tracking software (Noldus Information Technology, The Netherlands).

### Opto-RT-CPP

A three-compartment apparatus (Spatial Place Preference Box, Panlab, Harvard Apparatus) was used for the Opto-RT-CPP test. The apparatus is composed of two compartments with different visual and tactile cues for both floors and walls and a connecting corridor (neutral compartment) with transparent walls and floor. The study was carried out throughout 8 consecutive days preceded by one *“Habituation”* day. On each day of the experiment, subjects were placed in a neutral cage for at least 3 minutes to recover after connecting the optic cables. Mice were subsequently moved into the transparent corridor of the apparatus where, after 30-50 s, the doors were removed and animals allowed to freely explore the apparatus. The position of the mouse was detected by a camera positioned above the apparatus. On *“Habituation”*, *“Pre Test”* and “*Conditioning Place Test”* days (15 min), mice were connected to the optic fiber but no light was paired to any chamber. On the “*Real Time Conditioning”* days (30 min), one chamber was randomly chosen as a light-paired chamber (counterbalanced across animals) while the other one was subsequently assigned as light-paired chamber on the “*Reverse Real Time Conditioning”* days (30 min). Every time the subject entered the light-paired chamber, the laser was activated according to the stimulation protocol. Cumulative duration of the time spent in each compartment was recorded by using the Ethovision XT13.0/14.0 tracking software (Noldus Information Technology, The Netherlands). Mice that showed strong initial preference during the “Pre-Test” for either one of the two compartments (<25% or >75% of time spent) were excluded from the statistical analysis.

### Opto-Avoidance-EPM

The re-designed version of the EPM test aims to assess the aversive effect induced by the photostimulation in comparison to the natural aversion experienced in the open arms of the apparatus. In the Opto-Avoidance-EPM test, photostimulation was activated upon entry into any of the closed arms, and disabled by leaving it. In contrast, visiting the open arms or occupying the center zone had no effect on photostimulation. The animal was placed in the center zone of the arena and allowed to freely explore the apparatus for 15 min. Time spent in each compartment and number of entries were recorded by the Ethovision XT13.0/14.0 tracking software (Noldus Information Technology, The Netherlands).

### Opto-Negative Reinforcement paradigm (Opto-NR)

Mice response was assessed by using operant conditioning boxes equipped with two nose-poke devices (MED-PC, Med Associates inc, Fairfax, USA) and laser for optogenetic stimulation. One device was defined as active, which when activated terminates the optogenetic stimulation schedule, while the other one was demarcated as inactive, which had no control over the laser. The presentation of the optogenetic stimulations (10 s) were alternated with variable-time intervals, when no stimulation was applied. An active nose-poke (FR1) at any point during this pre-stimulation interval or during the stimulation epoch itself terminates the optogenetic stimulation schedule and produces a visible safety signal (house light ON) that marks a period of safety.

### Sugar Positive Reinforcement Paradigm (Sugar-PR)

Under this protocol mice learned to make active nosepokes to receive sugar pellets. The experiment is carried out during several consecutive sessions (30 minutes; 1 session/day). Mice were food restricted by administering one daily feeding of 2.5- 3g of standard food following each daily session.

Training and testing took place in operant boxes (MED-PC, Med Associates inc, Fairfax, USA) interfaced with lasers for optogenetic stimulation and equipped with nosepoke (NP) devices on each side of a food dispenser. One NP device was designated as active and marked by a light while the device on the other side of the dispenser was selected as inactive. Nose-poking to the active NP carried out when the light is on (“delivering phase”), names as “Active delivering NPs”, resulted in a cue tone presentation (0.5s), a 20 mg sucrose pellet delivery (5TUT, TestDiet, St. Louis, USA) and/or laser activation for optogenetic stimulation (20Hz, 5ms, 5mW) according to the different phases of the task, while nose-poking to the inactive side resulted in no cues presentation, sugar delivery and/or laser activation. During 20s that follows a sugar pellet delivery and/or optogenetic stimulation the visual cue for the active NP goes off (“no delivering phase”) and active nose-poking during this period did not result in auditory cue presentation, sugar delivery and/or laser activation. Active NPs carried out during this period were registered as “Active no delivering NPs” and summed to the “Active delivering NPs” to obtain the total Active NPs. Nose pokes in the inactive hole, did not activate the sugar dispenser, the laser and the acoustic cue at any time but they were registered as “Inactive delivering NPs” when carried out during the “delivering phase” and as “Inactive no delivering NPs” when performed during the “no delivering phase”. “Inactive no delivering NPs” were summed to the “Inactive delivering NPs” to obtain the total Inactive NPs.

Phase 1 (Acquisition phase): Sugar was delivered in response to an Active delivering NPs. This phase lasts until the animal acquires a stable sugar SA rate, which consists of at least 10 received sugar pellet/session with a <20% variation during three consecutive days. Animal that did not acquired a stable sugar SA rate within the third week were excluded from the experiment.

Phase 2 (Sugar-Stimulation phase): Optogenetic stimulation (10 s stimulation 20Hz, 5ms, 5mW) was delivered as a consequence of an active delivering NPs simultaneously to the activation of the sugar dispenser. This phase lasts 5 consecutive sessions (1 session/day).

Phase 3: Depending on the response upon pairing sugar delivering and optogenetic stimulation mice were exposed either to a “only sugar” or to an “only stimulation” phase 3. In the case of a reduction in the number of active NPs, and in order to exclude that such response was not due to a loss of interest for sugar, the subsequent and last phase is characterized “sugar reinstatement” during the following phase 3, restoring the same conditions previously applied during phase 1. If NP activity was similar or higher during phase 2 compared to phase 1 mice were moved to the “only stimulation” phase 3 (5 sessions) in order to assess whether optogenetic stimulation alone was sufficient to maintain a comparable NP activity.

Phase 4: Depending on the response upon “only stimulation” phase 3 mice were exposed either to a “only sugar” phase 4 or to an “only cues” phase 4. In the case of extinction of the active response, the subsequent and last phase 4 (2 sessions) was characterized by “sugar reinstatement” restoring the same conditions previously applied during phase 1. If NP activity was not affected during phase 3 compared to phase 2 mice were moved to the “only cue” phase 4 (3 sessions) where mice were only exposed to visual and auditory cues previously accompanied with sugar and optogenetic stimulation.

Phase 5: Response extinction was finally asses in a last phase (3 sessions) in which active NP activity was followed by no sugar delivery, no optogenetic stimulation and no cues presentation.

### Statistics

Data are expressed on the plots as means ± SEM. For behavioral analysis, repeated measures two-way ANOVA were performed followed by Šídák or Tukey multiple comparisons where appropriate. For single cell extracellular recordings, Friedman test was performed followed by Dunn’s multiple comparisons. For *ex vivo* electrophysiology, paired t-test was used (GraphPad Prism version 7.00 for Windows, GraphPad Software, La Jolla California USA). See Supplemental file 1.

### Post-injection histological analysis

Following behavioral analyses, recombinase mice were deeply anesthetized and perfused trans-cardially with phosphate-buffer-saline (PBS) followed by ice-cold 4% formaldehyde. Brains were extracted and 60 μm sections were cut with a vibratome. Fluorescent immunohistochemistry was performed to enhance the YFP signal. All mice were analysed. Mice that displayed strong cellular YFP labeling in the expected area (STN or pSTN) and in which optic cannulas could be confirmed as positioned immediately above the STN or pSTN, respectively were included in the statistical analyses of the electrophysiological and behavioral experiments.

After rinsing in PBS, sections were incubated for 3 hours in PBS containing 0.3% X- 100 Triton and 5% blocking solution (normal donkey serum) followed by incubation with primary antibodies diluted in 5% normal donkey serum in PBS, overnight at 4 ◦C (chicken anti-GFP 1:1000, cat. no. ab13970, Abcam; rabbit anti-c-Fos 1:800, cat. no. 226003, Synaptic Systems; rabbit anti-c-Fos 1:800, cat. no. ab190289, Abcam). Next day, sections were rinsed in PBS and incubated for 2 hours with secondary antibodies diluted in PBS containing 0.3% X-100 Triton and 5% normal donkey serum (A488 donkey anti-chicken 1:1000, cat. no. 703–545-155, Jackson ImmunoResearch; A594 donkey anti-rabbit 1:1000, cat. no. ab150068; Abcam). After rinsing in PBS, sections were incubated for 10 min with DAPI diluted in distilled water (1:5000). Sections were mounted with Fluoromount-G® mounting medium (SouthernBiotech) and cover- slipped. Analysis was performed upon slide scanning using the NanoZoomer 2–0- HT.0 (Hamamatsu) scanner followed by visualization using the NDPView2 software (Hamamatsu).

For enzymatic immunohistochemistry mice were deeply anesthetized and perfused trans-cardially with phosphate-buffered-saline (PBS) followed by ice-cold 4% formaldehyde. Brains were extracted and 40 μm and 60 μm sections were cut with a cryostat and a vibratome, respectively. After rinsing in PBS, the endogenous peroxidase inhibition was performed in PBS containing 0.3% X-100 Triton and 1% H_2_O_2_ for 15 min. Sections were rinsed in PBS and incubated for 90 min in PBS containing 0.3% X-100 Triton and 5% blocking solution (normal goat serum) followed by incubation with primary antibody diluted in 5% normal goat serum in PBS, overnight at 4 ^◦^C (rabbit anti-c-Fos 1:800, cat. no. 226003, Synaptic System; rabbit anti-c-Fos 1:800, cat. no. ab190289, Abcam). Next day, sections were rinsed in PBS and incubated for 90 min in a solution of biotinylated goat anti-rabbit IgG (Vector Laboratories®) at a dilution of 1:500 in PBS containing 0.3% X-100 Triton and 5% normal goat serum. After rinsing in PBS, sections were incubated for 90 min with the mixed avidin–biotin horseradish peroxidase (HRP) complex solution with PBS (ABC Elite Kit, Vector Laboratories®). Sections were rinsed in PBS and washed for 10 min with 0.1 M Tris-HCl at pH 7.4. The peroxidase complex was visualized by exposure to a chromogen solution containing 3,3′-diaminobenzidine tetrahydrochloride (DAB, Vector Laboratories®) in distilled water. The reaction was stopped by washing in 0.1 M Tris-HCl at pH 7.4. After rinsing in PBS, sections were mounted, air-dried, and stained in cresyl violet for nuclear staining. Then, sections were dehydrated in increasing concentrations of ethanol and cleared in Histolab Clear®. Finally, the coverslips were glued to the slides using DePeX mounting media. Analysis was performed upon slide scanning using the NanoZoomer 2–0-HT.0 (Hamamatsu) scanner followed by visualization using the NDPView2 software (Hamamatsu).

